# The proofreading mechanism of the human leading strand DNA polymerase ε holoenzyme

**DOI:** 10.1101/2025.04.11.648458

**Authors:** Feng Wang, Qing He, Michael E O’Donnell, Huilin Li

## Abstract

**Significance Statement:** Replicative DNA polymerases (Pol) function with a sliding clamp and their proofreading exonuclease provides high fidelity. Thus, study of the proofreading mechanism must both contain the Pol-clamp complex and generate the mismatch in situ to ensure the mismatched primer follows a physiological route from the pol to exo site. Despite numerous previous studies, none satisfy both criteria. This study on human Polε–PCNA meets both criteria. Cryo-EM analysis captured proofreading intermediates that reveal an unexpected proofreading process. The primer is unwound by 6 nucleotides that scrunches to form out-of-register base pairs with the template. These findings provide new insights and calls for reevaluation of proofreading mechanisms performed without a sliding clamp or with an existing mismatch.

The eukaryotic leading strand DNA polymerase epsilon is a dual function enzyme with a proofreading exonuclease site located 40 angstroms from the DNA synthesizing polymerase site. Errors in Pol epsilon proofreading can cause various mutations, including C to G transversions, the most prevalent mutation in cancers and genetic diseases. Pol epsilon interacts with all three subunits of the PCNA ring to assemble a functional holoenzyme. Despite previous studies on proofreading of several polymerases, how Pol epsilon, or any Pol complexed with its sliding clamp proofreads a mismatch generated in situ has been unknown. We show here by cryo-EM that a template/primer DNA substrate with a pre-existing mismatch cannot enter the exo site of Pol epsilon/PCNA holoenzyme, but a mismatch generated in the Pol site yields three proofreading intermediates of Pol epsilon/PCNA holoenzyme. These intermediates reveal how the mismatch is dislodged from the Pol site, how the DNA unwinds 6 base pairs and how the unpaired primer 3′-end is inserted into the exo site for cleavage. These results unexpectedly demonstrate that PCNA imposes strong steric constraints that extend unwinding and direct the trajectory of mismatched DNA, and that this trajectory is dramatically different than for Pol epsilon in the absence of PCNA. These findings suggest a physiologically relevant proofreading mechanism for the human Pol epsilon/PCNA holoenzyme.

## Introduction

DNA replication is a fundamental process required of all cell types (1–3). Replicative polymerases, such as the leading strand Polε and lagging strand Polδ synthesize DNA with extraordinarily high fidelity (4–7). The high accuracy is achieved by their 3′-5′ exonuclease activity (8–10), which correct errors when they are made by the DNA polymerase (pol) (**Fig. 1A**). Human Polε belongs to the B-family of DNA Pols and contains 4 subunits, the large catalytic subunit POLE1 (Pol2 in yeast) and three regulatory subunits POLE2-4 (Dbp2-4 in yeast) (11, 12). *POLE1* has undergone a gene fusion between two distinct but related B family DNA polymerases (13, 14). The fusion protein contains a catalytic N-terminal domain (NTD) and a noncatalytic C-terminal domain (CTD) (**Fig. 1B**); the NTD and CTD are linked by a long loop that interacts with the two small histone-fold POLE3 and POLE4 subunits (15). The CTD is unique to POLE1 (16, 17), and together with POLE2, forms a complex with the replicative helicase Cdc45-Mcm2-7-GINS (CMG) to produce the leading strand replisome (18–21). The POLE1 NTD harbors both DNA polymerase and 3′-5′ proofreading exonuclease (Exo) activities(22, 23), and is therefore referred to as Polε-core in this study.

**Fig. 1.**
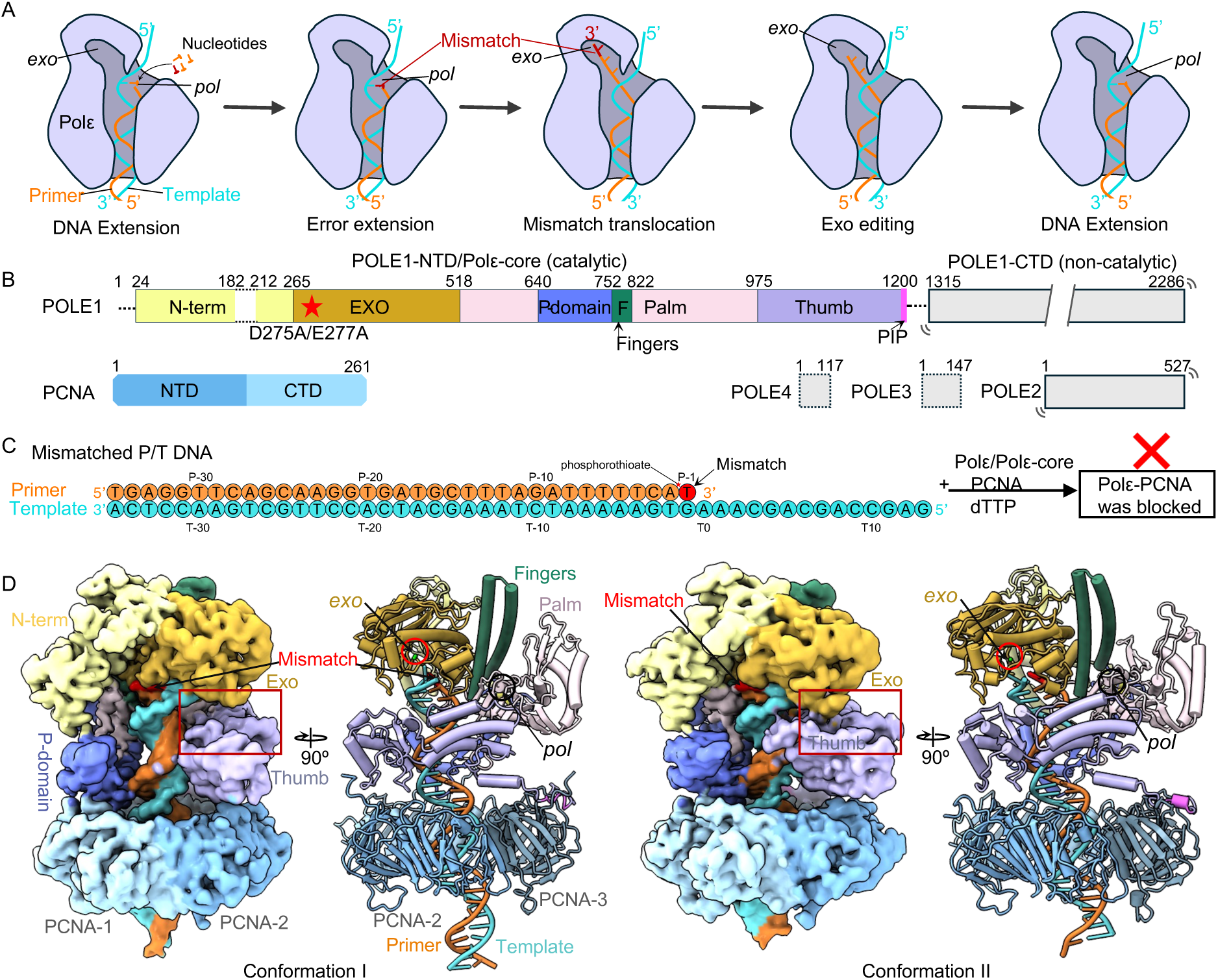
Cryo-EM structures of Polε–PCNA bound to T/P with a pre-existing mismatch. (A) sketch of the Polε holoenzyme proofreading process based on published studies in the absence of sliding clamp. The T/P unwinds by 3 bp and the unwound regions are kept separated during mismatch editing. The *pol* and *exo* sites are labeled. (B) Domain architectures of PolE1-4 and PCNA. Dashed lines indicate disordered regions. The structurally resolved regions (POLE1-NTD, i.e., Polε-core, and PCNA are colored by domains. POLE1-CTD and POLE2-4 are mobile relative to POLE1-NTD and invisible in EM maps. (C) The nucleotide sequence of the primer and template DNA. The mismatched primer 3’-end is highlighted in red. The T/P with a pre-existing mismatch cannot reach the *exo* site of Polε holoenzyme and binds Polε in a blocked state. (D) EM maps and atomic models of the Polε-PCNA–T/P complex in two blocked conformations. The maps and models are labeled and colored by domains as in (B). The distinct interactions of the Exo and thumb domains in the two conformations are highlighted by the red box.

The Polε-core contains the conserved thumb, fingers, and exo domains plus a unique P-domain (23) (**Fig. 1B**). Polε-core interacts with the ring-shaped PCNA sliding clamp to assemble the holoenzyme. We and others recently solved the structures of the Polε-core–PCNA–DNA ternary complex in the DNA polymerizing state, revealing that Polε-core contacts all three subunits of the PCNA trimer via its C-terminal PIP motif, thumb domain, and P-domain, respectively, and that the P-domain contributes to Polε processivity by binding to both PCNA and dsDNA (24–26). These structures combined with previous activity assays underscore the importance of the PCNA sliding clamp in Polε activity (27, 28), and are consistent with the functional requirement of a sliding clamp of all other replicative polymerases (29, 30). While the canonical RFC PCNA clamp loader can load PCNA onto either strand (31–33), we recently discovered that PCNA loading by the leading strand specific CTF18-RFC loader is facilitated by the Polε P-domain by stabilizing CTF18-RFC for PCNA opening and DNA loading (34).

Proofreading occurs after the DNA Pol incorporates a mismatched nucleotide in the *pol* site, followed by unwinding a short segment of template/primer (T/P) to enable the 3’ end of the primer to relocate at a distant *exo* site for mismatch removal. Single-molecule analysis showed that Pol alone, without its sliding clamp, can rapidly dissociate from the mismatched DNA at the pol site and rebind DNA at its exo site during in vitro replication (35, 36), but a dissociation-and-reassociation mechanism is not expected for a processive Pol–clamp complex that binds DNA without dissociation (i.e. processive action). In other words, to capture authentic proofreading intermediates as expected to occur in vivo, an in vitro system must minimally ensure that the Pol is attached to its sliding clamp, from misincorporation in the Pol site to excision in the exo site. We find here that Polε misincorporation and proofreading can occur within the Polε–PCNA complex. Interestingly the processive action of Polε confined by PCNA-DNA imposes an unexpected, but important constraint on how the enzyme and DNA move during proofreading.

To ensure that the mismatch in fact occurs at the *pol* site before the mismatch enters the *exo* site, the mismatch should be generated in situ by allowing mismatch formation to proceed to proofreading. If instead one were to provide a preformed mismatch DNA to Polε, it will directly bind the *exo* site without first going through the *pol* site, and the result may not reflect the in vivo situation in which the mismatch originates in the Pol site. This is important because the spatial and temporal coupled processes of the T/P unwinding and *pol*-to-*exo* translocation might be expected to occur in the cell as a processive Polε–PCNA complex.

Several DNA Pol structures, without their cognate clamp, have been reported in the proofreading states, including *E. coli* Pol I (37), *E. coli* Pol III (38), bacteriophage RB69 Pol (39–41), archaeal PolB (42), mitochondrial Polγ (43, 44), and human Polε (25). These structures have suggested a largely conserved proofreading mechanism in which a mis-incorporated nucleotide at the primer 3’ end fails to pass the Watson-Crick base-pairing checkpoint in the *pol* site and is blocked from further polymerization (**Fig. 1A**). This triggers Pol backtracking, primer unwinding 3 base pairs (bp) from the template, and primer translocation to the *exo* site, where the mismatch is removed (45, 46). The excised primer then translocates back to the *pol* site to resume DNA synthesis. The *E. coli* replicase, DNA Pol IIIα (with its clamp), was also found to unwind 3 bp of the T/P (47), and archaeal PolD-PCNA unwinds 5 bp (48). But all these studies used either Pol in the absence of a sliding clamp or used pre-synthesized mismatch DNA in which a clamp may not encircle DNA during the entire process of mismatch formation and excision. In a recent study of human Polε proofreading, the in vitro system lacked a stably bound PCNA and used a preexisting DNA mismatch (25). Considering Pol proofreading studies thus far lacked a stably bound PCNA, the proofreading mechanism of a processive DNA Pol-clamp complex is not yet known and is a goal of the present study.

In the current study, we use a sufficiently long DNA to stabilize the PCNA clamp and generate the mismatch in situ. In the presence of the PCNA clamp, we find unique results not observed before for any Pol, including Polε lacking PCNA. We first demonstrate by cryo-EM that a T/P with a preexisting mismatch is blocked from entering either the *pol* site or the *exo* site of the human Polε–PCNA holoenzyme. We capture three authentic proofreading intermediates only when a mismatch is first produced in situ at the *pol* site of human Polε–PCNA holoenzyme. These intermediates represent the mismatch-locking state, Pol-backtracking state, and mismatch-editing state. In contrast to the 3-bp melting by the conserved B-family DNA Pol alone or Polε in the absence of PCNA (10, 25), we observe 6 bp are melted by the processive Polε–PCNA. In all three intermediate states, Polε-core is stably associated with PCNA similar to the DNA synthesizing mode. This study is the first in which proofreading intermediates are captured in the context of a functional holoenzyme, and the observed movement of mismatched primer 3’-end is actually originated from the *pol* site. We therefore suggest that the proofreading mechanism revealed by these intermediates may be the physiologically relevant process inside the cell.

## Results

### 1. Polε holoenzyme with a pre-existing mismatch could not enter a proofreading state

A T/P DNA substrate with a 20-23 bp dsDNA region is sufficient to span the human Polε and PCNA in the polymerizing state, but not long enough to maintain stable PCNA binding in the proofreading state (24, 25), because Polε backtracks and unwinds several bp during proofreading (25). We therefore designed T/P substrates with a 35-bp dsDNA region to investigate the proofreading mechanism by the human Polε–PCNA holoenzyme (**Fig. 1C**). We first asked if a T/P containing a pre-existing mismatch (G•T) can form the expected proofreading intermediates by the Polε–PCNA holoenzyme. We mixed the purified human Polε, PCNA, and the mismatched T/P DNA at a molar ratio of 1:3:1.1 in the presence of 0.5 mM dTTP at room temperature for 10 minutes and incubated the mixture on ice for 2 hours prior to preparing cryo-EM grids (***SI Appendix,* Fig. S1A**). Cryo-EM imaging and associated 2D and 3D classifications and 3D reconstruction resulted in two 3D EM maps of the Polε–PCNA–DNA ternary complex at an average resolution of 3.88 Å and 3.81 Å, respectively (**Fig. 1D, *SI Appendix,* Fig. S1 and S2**).

Atomic modeling of the two maps shows that Polε-core stably binds to the PCNA and P/T, but the POLE1-CTD and the regulator subunits POLE2-4 are invisible (***SI Appendix,* Fig. S1B-H**), similar to Polε holoenzyme in the polymerizing state (24, 25). This is consistent with the knowledge that the two Pol lobes of Polε are flexibly linked(19–21). Notably, the template strand extends to the exo site in both structures (**Fig. 1D, *SI Appendix,* Fig. S2**), indicating that the T/P with a pre-existing mismatch cannot reach the proofreading state in the Polε holoenzyme when the DNA movement is highly constrained by PCNA. Instead, the T/P junction binds just below the Exo and N-term domains in the highly positively charged main substrate chamber (***SI Appendix,* Fig. S2**), thus placing the holoenzyme in a blocked state.

The Exo domain is distant from the thumb in one conformation but interacts with the thumb via a thumb-domain-contacting loop (TCL) in the other conformation (***SI Appendix,* Fig. S1A-B**). Therefore, the template strand passes through a gap between the TCL and the thumb domain in the first conformation but is blocked by the TCL in the second conformation. While the upper end of the T/P DNA substrate is blocked by Polε, the bottom dsDNA end of the T/P protrudes from PCNA by 6 bp (**Fig. 1D**). As described below, in the authentic proofreading state, no stable dsDNA density is observed below the PCNA, as the T/P moves upwards and is unwound for proofreading.

### 2. Polε enters an authentic proofreading state upon encountering an in-situ mismatch

We next designed a T/P substrate lacking a mismatch. The DNA substrate contains a 29-bp duplex region to ensure stable PCNA binding during Polε proofreading and an 18-nt tail in the template strand (AAAGT-GAAAC-GACGA-CCG) (**Fig. 2A**). In the presence of dTTP, the primer will be extended by three nucleotides (3 T’s) by Pol, followed by a G•T mismatch due to the lack of dCTP. Therefore, this experimental design results in a mismatch that is generated in situ within the *pol* site (44, 49), ensuring that the subsequent movement of the mismatch actually originates from the *pol* site, in a manner that mimics in vivo proofreading.

**Fig. 2.**
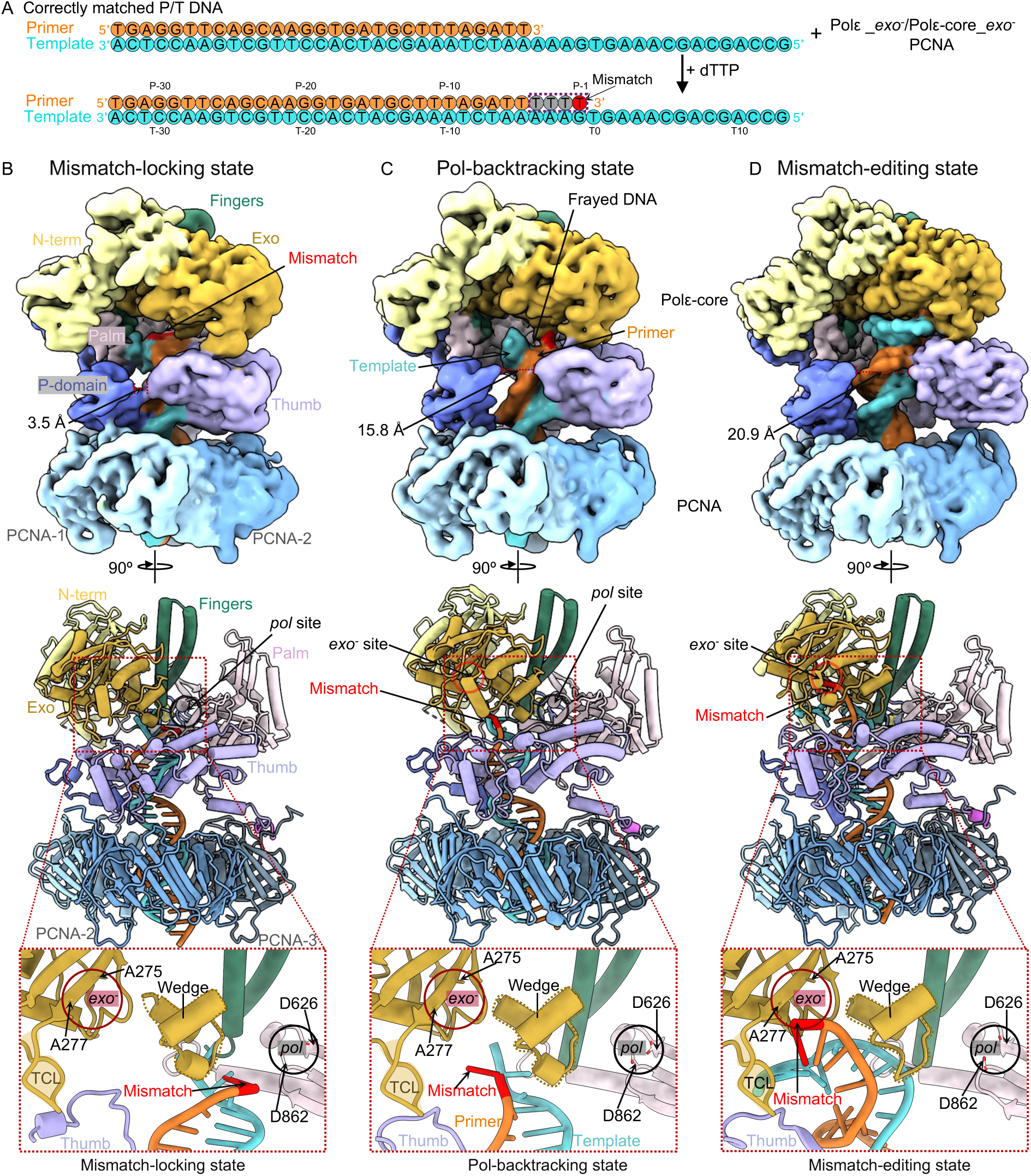
Cryo-EM structures of Polε–PCNA bound to an in-situ generated mismatch. (A) Scheme for capturing Polε holoenzyme in the proofreading states. Nucleotide sequence of the P/T is shown in color. In the presence of dTTP, Polε extends 4 T; 3 are matched with template (gray) and the last one is mismatched (red). (B-D), Cryo-EM maps and atomic models of the ternary complex in the mismatch-locking (B), Pol-backtracking (C), and mismatch-editing (D) states. Cryo-EM maps are shown in the upper panel, atomic models in the middle, and close-up views of the mismatched primer 3’-end in the three proofreading intermediates in the lower panel.

We initially pre-incubated a Polε Exo^-^ variant (D275A/E277A(50); **Fig. 2A**), PCNA, and the above-described P/T DNA substrate at a molar ratio of 1:3:1.1, then added 0.5 mM dTTP to the reaction and incubated the mixture at room temperature for 3 minutes prior to making cryo-EM grids. But the assembled complex particles had preferred orientation issues that prevented us from obtaining a good 3D map. Considering that only the catalytic Polε-core binds to PCNA (24, 25), we substituted Polε-core Exo^-^ in place of full length Polε (we note that the proofreading activity of Polε-core is comparable to that of full-length Polε (51).

The in vitro assembled complexes were heterogeneous, as expected for different stages of the proofreading process. By 3D classification and 3D reconstruction, we obtained three 3D EM maps of the Polε-core–PCNA–DNA ternary complex at an average resolution of 3.60 Å, 3.53 Å, and 3.11 Å resolution, respectively (**Fig. 2B-D, *SI Appendix,* Fig. S3**). This process provided three atomic models (**Fig. 2B-D, *SI Appendix,* Fig. S4**). In all three structures, the Polε-core fingers domain is in the open conformation such that the Pol activity is arrested, and the primer 3′-end is being transferred from the *pol* to *exo* site, clearly indicating that these states represent proofreading intermediates of the holoenzyme. Importantly, Polε-core stably binds to PCNA via its P-domain, thumb domain, and the C-terminal PIP motif (**Fig. 2B-D**), in a manner that largely resembles the PCNA interaction when the holoenzyme is in the DNA polymerizing states(24, 25). Therefore, these new structures clearly show that PCNA remains stably associated with Polε and will strongly constrain T/P movement due to PCNA encirclement of DNA. This also limits the movement of the 3’ terminal mismatch relative to the enzyme. We therefore observe very different proofreading behavior of the holoenzyme compared to a previous study performed in the absence of PCNA (25), as detailed below.

### 3. Three proofreading intermediates of the human Polε

Here we describe the DNA binding mode in each of the three proofreading states, with a focus on the location of the in situ synthesized primer 3′-end mismatch.

The first intermediate is a “mismatch-locking state”. The overall Polε-PCNA structure in this state is nearly identical to the polymerizing state of Polε-PCNA (24, 25), except the fingers switch from a closed to an open conformation (***SI Appendix,* Fig. S5A-B**). Consistently, the pol site in the first intermediate lacks an incoming nucleotide (**Fig. 2B**), and the 3′ mismatched primer end is 1 bp below the post-insertion site of Polε (***SI Appendix,* Fig. S5C**). Previous studies reveal that the incoming nucleotide cannot be incorporated into the 3′ mismatched primer end due to a failure of Watson-Crick base checking in the post insertion site of DNA Pol (52–54). Thus, when the post-insertion site adopts a 3′ mismatch base, the polymerizing state could mimic the mismatch sensing state. Compared to these states, the holoenzyme has moved forward by 1 bp while maintaining tight binding to the T/P in the first state (**Fig. 2B-C, *SI Appendix,* Fig. S5C**). Consequently, the mismatch is locked 1 nucleotide below the pol site and cannot be extended by Pol activity. For this reason, we have referred to the first state as the mismatch-locking state in which the *pol*-to-*exo* transition has initiated. Interestingly, when Polγ (a family D Pol) synthesized a mismatch in situ, it also moved forward by 1 bp after mismatch sensing (44).

The second intermediate is in a Pol-backtracking state. In this state, the EM density of the T/P DNA substrate is weaker and frayed such that the unwinding has started (**Fig. 2C**). Because the unwound primer 3′ mismatch is at the entry of the channel leading towards the *exo* site (**Fig. 2C**), this state represents the Polε-backtracking state in which initial melting has allowed the primer 3′-end to move away from the pol and toward the exo site. Further, the thumb, along with the dsDNA region, has moved away from the P-domain to facilitate the T/P fraying (**Fig. 2B-C**).

The third intermediate is in a mismatch-editing state. The T/P DNA substrate has stronger EM density in this state than in the first and second states, indicating it is a relatively stable state of the holoenzyme (**Fig. 2D**). The mismatched primer 3′-end has fully translocated into the *exo* site (**Fig. 2D**). In a wild-type Polε, the mismatch would have been cleaved in this state (8), but because we used the mutant Polε exo^-^ in this study there is no cleavage.

### 4. Mismatch translocation from the mismatch-locking to Pol-backtracking state

Superimposition of the Polε-core backtracking and mismatch-locking states reveals that the P-domain and the α/β subdomain of the thumb moves away from the dsDNA region by 3.6 Å and 6.8 Å, respectively (**Fig. 3A**). The larger space between the P-domain and the thumb allows a longitudinal movement of the T/P by 1 bp in the backtracking state. The PCNA undergoes a rigid-body movement, but the interaction between the P-domain and PCNA-1, and between the PIP motif with the extended β-strand and PCNA-3, remain unchanged (**Fig. 3B**). These interactions resemble the corresponding interactions between Polε-core and PCNA in the polymerizing state(24, 25) (***SI Appendix,* Fig. S5D**). However, the α/β subdomain of the thumb, which only interacts with PCNA very weakly, moves outwards by ∼ 8-9 Å above PCNA-2. And this leads to a 2-3 Å outward movement of the helix-bundle subdomain of the thumb, which in turn, results in a ∼10° outward rotation of the linker helix connecting the PIP motif and the helix-bundle subdomain (**Fig. 3B**). Overall, the three-point interaction between Polε-core and PCNA is largely sustained transitioning from the mismatch-locking to Pol-backtracking state.

**Fig. 3.**
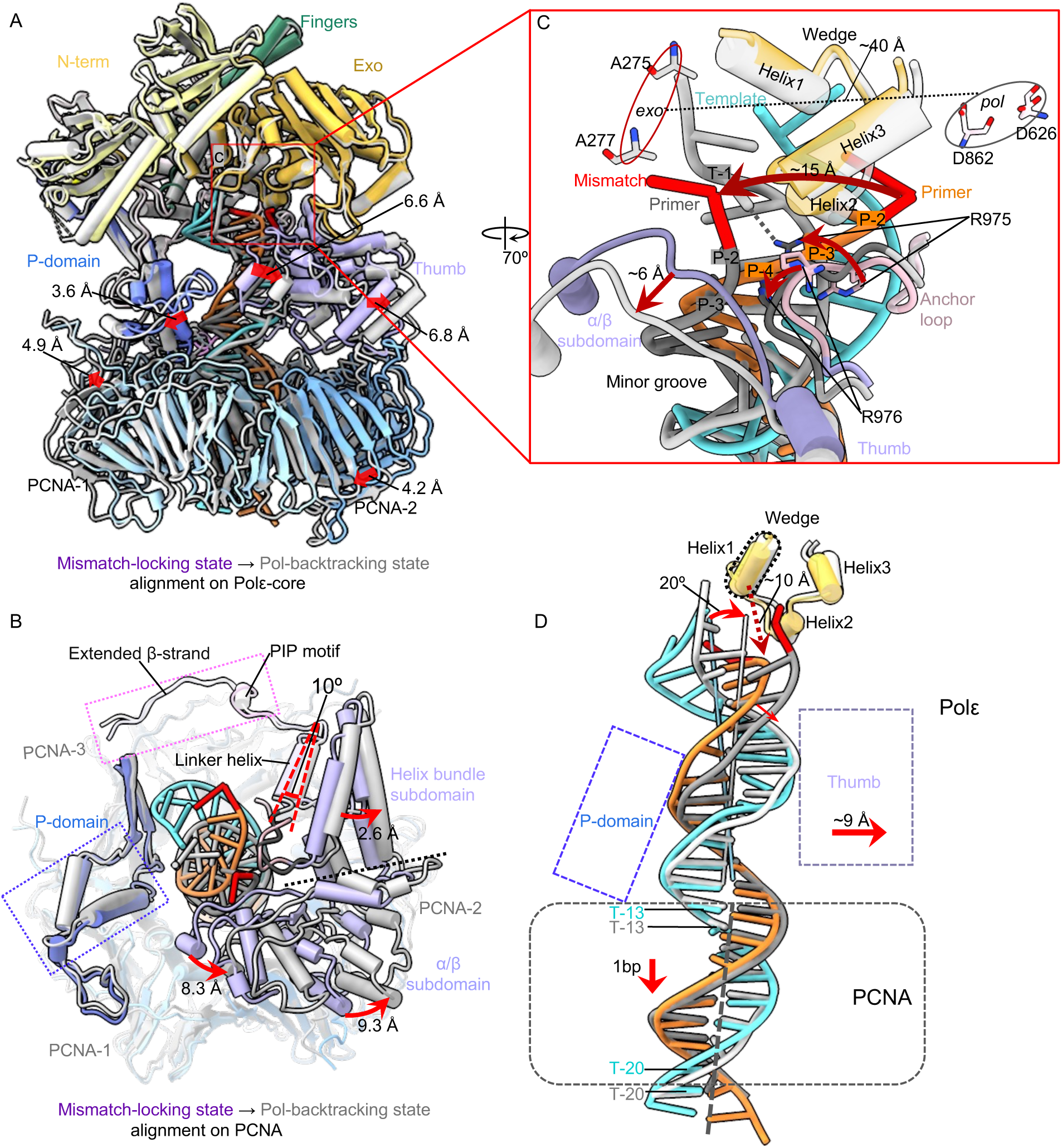
Conformational changes of the T/P from the mismatch-locking to Pol-backtracking state. (A) Superimposition on Polε-core of the mismatch-locking (color) and Pol-backtracking states (gray). Domain movements are indicated by red arrows. (B) A top view of the superimposition at the Polε-PCNA interface where most movements occur. The bulk of Polε-core is omitted for clarity. (C) Close-up view of changes in the mismatched region of T/P between the two states. The movements are indicated by red arrows and labeled. (D) Superimposition of T/P based on alignment on PCNA in both states. The DNA tilts 20° and moves with the thumb outwards by 9 Å. And the DNA moves down by 1 bp.

The *pol* and *exo* sites are ∼40 Å apart (**Fig. 3C**). The mismatched primer 3′-end moves about 15 Å from the locking to backtracking state, and the primer ends in these two states are separated by a three-helix wedge in the Exo domain. It appears that the inner α/β subdomain of the thumb has pushed the primer at the P-2/P-3 region to a distinctive position in the Pol-backtracking state. Further, the loop linking the palm and thumb domains contains multiple positively charged residues that may anchor the T/P transition during proofreading (25, 51). For example, Arg-975 and Arg-976 seem to reposition the primer at the P-3/P-4 region, and both residues are inserted in the T/P minor groove (**Fig. 3C**). Importantly, Arg-975 appears to directly contribute to the T/P fraying by forming a H-bond with the template T-1 base, which enables the mismatched primer 3’-end to flip toward the *exo* site in the Pol-backtracking state. The dsDNA region inside PCNA slides down by 1 bp from the mismatch-locking to the backtracking state (**Fig. 3D**), and the dsDNA region inside the Polε-core tilts 20° toward the thumb domain, which moves away by ∼9 Å to accommodate the new DNA location. The wedge helix1 is 10 Å above the frayed T/P junction and is likely not responsible for the T/P separation, in contrast to the suggested Polε proofreading model (25). Overall, the thumb and the anchor loop drive the T/P transition from the locking to backtracking state.

### 5. Polε holoenzyme in the mismatch-editing state

The primer 3′-end reaches the *exo* site in the mismatch-editing state (**Fig. 4A**). The upper region of the T/P has well-defined EM density that resolves individual bases (**Fig. 4B**). There are three separated primer bases in the *exo* channel, similar to previous studies on B-family DNA Pols (25, 40, 55, 56) (**Fig. 4C-D**). The phosphate of the 3′-end P-1:dT faces a highly negatively charged region of the exo site, which would otherwise be mediated by two cations in the wild type exo site. But the 3′ dT ribose hydroxyl H-bonds with Met-444, and the P-1:dT base form a hydrophobic interaction with Pro-286 and Pro-441 (**Fig. 4C-D**). These interactions may be sufficient to stabilize the primer 3′-end in the mutated *exo* site (*exo^-^*). Superimposing the wild type exo site (PDB ID 9F6L(25)) with the current exo^-^ site reveals that the two catalytic Mg²⁺ ions align precisely to coordinate the phosphate of the mismatched P-1:dT (**Fig. 4D**). The following P-2:dT/P-3:dT region is stabilized by two H-bonds, with their respective phosphate H-bonds with Leu-424 and Gly-420, respectively (**Fig. 4D**).

**Fig. 4.**
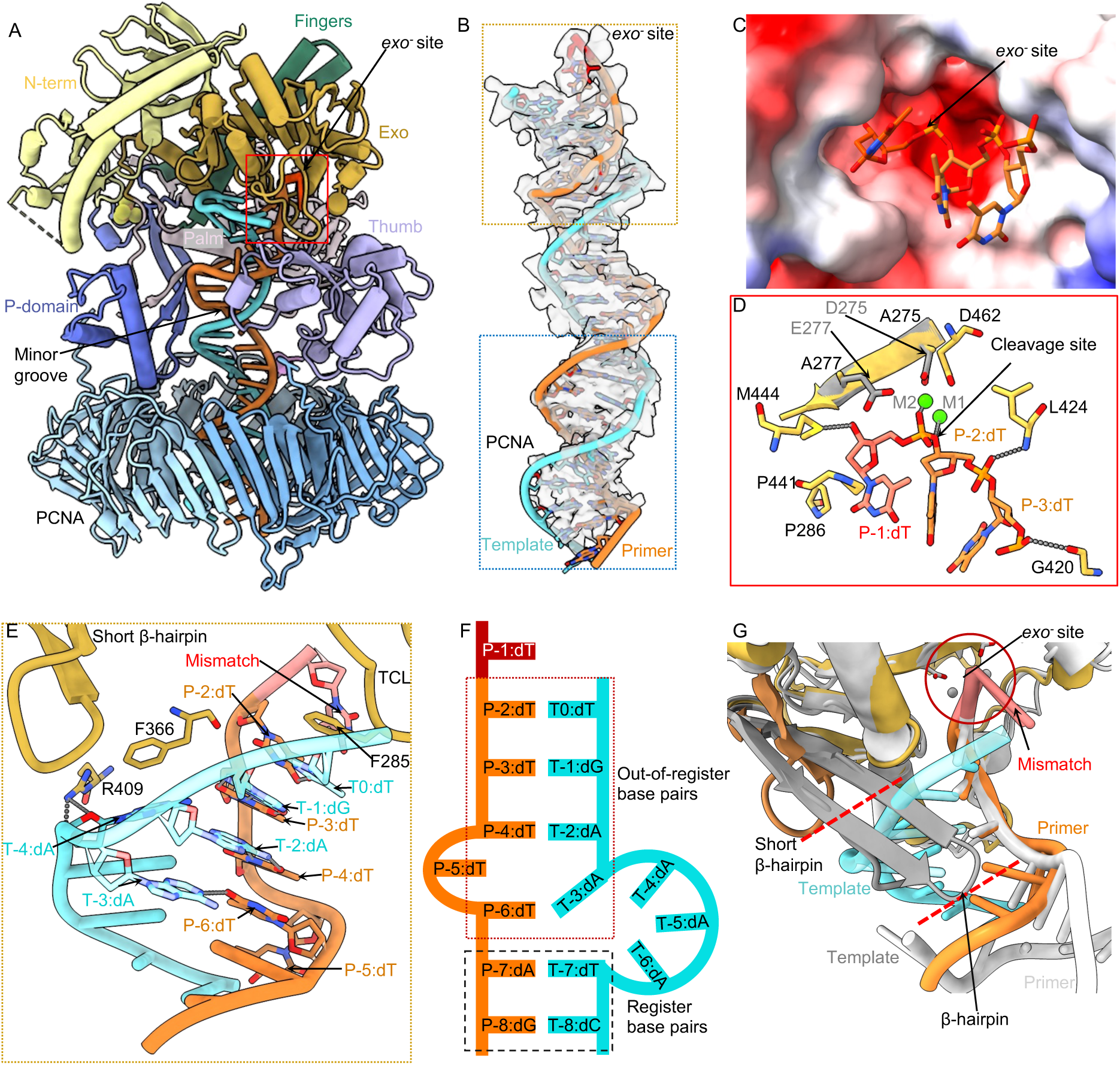
Polε interactions with T/P DNA in the mismatch-editing state. (A) The mismatch-editing state structure colored by domains. (B) T/P DNA in sticks superimposed with the EM density in transparent surface view. The top orange box shows the clear DNA density around the *exo^-^* site of Polε, the bottom blue box shows that the weak DNA density inside the PCNA clamp. (C) Electrostatic surface of the *exo^-^* site with the primer 3′-end. (D) The Polε *exo^-^* site structure. The catalytic D275 and E277 coordinating two Mg^2+^ in wild-type Polε are shown; they are mutated to A275 and A277 in Polε-exo^-^. (E) Close-up view of T/P around the *exo^-^* site. The T/P melts 6 bp then forms 4 out-of-register base pairs. The β-hairpin does not contact the template. Residues stabilizing the T/P are in sticks and labeled. (F) Sketch of T/P in editing state. (G) Comparison of the short Polɛ β-hairpin of (dark orange) and the longer RB69 gp43 β-hairpin (gray). The longer phage β-hairpin separates the template from primer.

A total of six bp are melted from the primer 3′-end in the editing state of the holoenzyme, the longest unwound stretch observed to date in any proofreading study (**Fig. 4E-F**). Unexpectedly, the template strand is scrunched by 2-3 nucleotides, such that the frame-shifted and melted template and primer form two mismatched base pairs (P-2:dT with T0:dT, P-3:dT with T-1:dG) and two serendipitously correct base pairs (P-4:dT with T-2:dA and P-6:dT with T-3:dA). But P-5:dT does not base-pair with the template (**Fig. 4E-F**). Therefore, the P-2 and P-3 bases in the exo channel are protected by the out-of-register base pairing, allowing only the mismatched primer 3’-end to be cleaved. The Polε exo domain adopts the DnaQ-like exonuclease fold conserved among A, B, and C-family DNA Pols(57). While all B-family DNA Pols possess a β-hairpin loop in the exo domain to facilitate T/P translocation between the *pol* and *exo* sites(55, 58, 59), the β-hairpin loop is much shorter in Polε (51) (**Fig. 4G**). The structure of the mismatch-editing state reveals that the β-hairpin is too short to interact with the template and to keep it separated from the primer (**Fig. 4E,G**). Instead, the template is only stabilized at T-4:dA by the Exo domain Arg-409 forming two H-bonds and the aromatic Phe-366 via a π-π interaction (**Fig. 4E**). We therefore suggest that the uniquely short β-hairpin loop underlies the out-of-register base-(mis)pairing in the T/P. Importantly, it is Phe-285 of the exo domain TCL that stabilizes the T0:dT via a π-π interaction to keep the template strand separated from the primer 3′-end (**Fig. 4E**). This perhaps explains why the neighboring residue Pro-286 is so important to the proofreading activity and that the P286K/R mutations are often found in cancers (50, 60, 61).

Beyond the exo domain, the Polε-core P-domain, fingers, palm, and thumb domains primarily interact with the dsDNA region (***SI Appendix,* Fig. S6**). Specifically, the P-domain Arg-672, Glu-674, Lys-733, and His-735, the fingers domain Lys-822, the palm domain Lys-950 and Lys-953, and the anchor loop Lys-974 and Arg-976 all interact with the duplex. In the thumb α/β subdomain, Ser-1038 engages the phosphate, and Ser-1040 H-bonds with thymidine base of P-4:dT (***SI Appendix,* Fig. S6A**). The DNA duplex is slightly tilted towards PCNA-1 in the PCNA chamber and is surrounded by positively charged residues such as PCNA-1 Lys-14, Lys-20, Lys-77, Arg-149, and Lys-217, PCNA-2 Lys-80 and Arg-149, and PCNA-3 Lys-80 and His-153 (***SI Appendix,* Fig. S6B**). But these residues are 4-6 Å away from the duplex phosphate backbones to allow DNA mobility, likely accounting for the weaker DNA density inside the PCNA ring (**Fig. 4B**).

### 6. Changes in Polε during transition from the Pol-backtracking to mismatch-editing state

The T/P undergoes dramatic transformation, as only one base at the mismatched primer 3′-end is frayed in the Pol-backtracking state, but six base pairs are melted from the primer 3′-end and rearranged in the mismatch-editing state (**Figs. 3-4**). Comparison of these two states reveals that the top region of Polε-core is well aligned but its PCNA contact region undergoes substantial conformational changes **(Fig. 5A).** Specifically, the palm domain loop moves (from orange and blue) to engages DNA with its Lys-950 and Arg-953, after transition to the mismatch editing state (**Fig. 5B**). And the anchor loop (primarily Arg-975) that originally inserted in the DNA minor groove (between white and gray), now flips upward and dissociates from DNA (between orange and blue) in the mismatch editing state (**Fig. 5B**). Because the local DNA movement directions coincide with the movement directions of both the palm and anchor loops as described above, we suggest that the T/P transformation is driven by these two Polε-core loops.

**Fig. 5.**
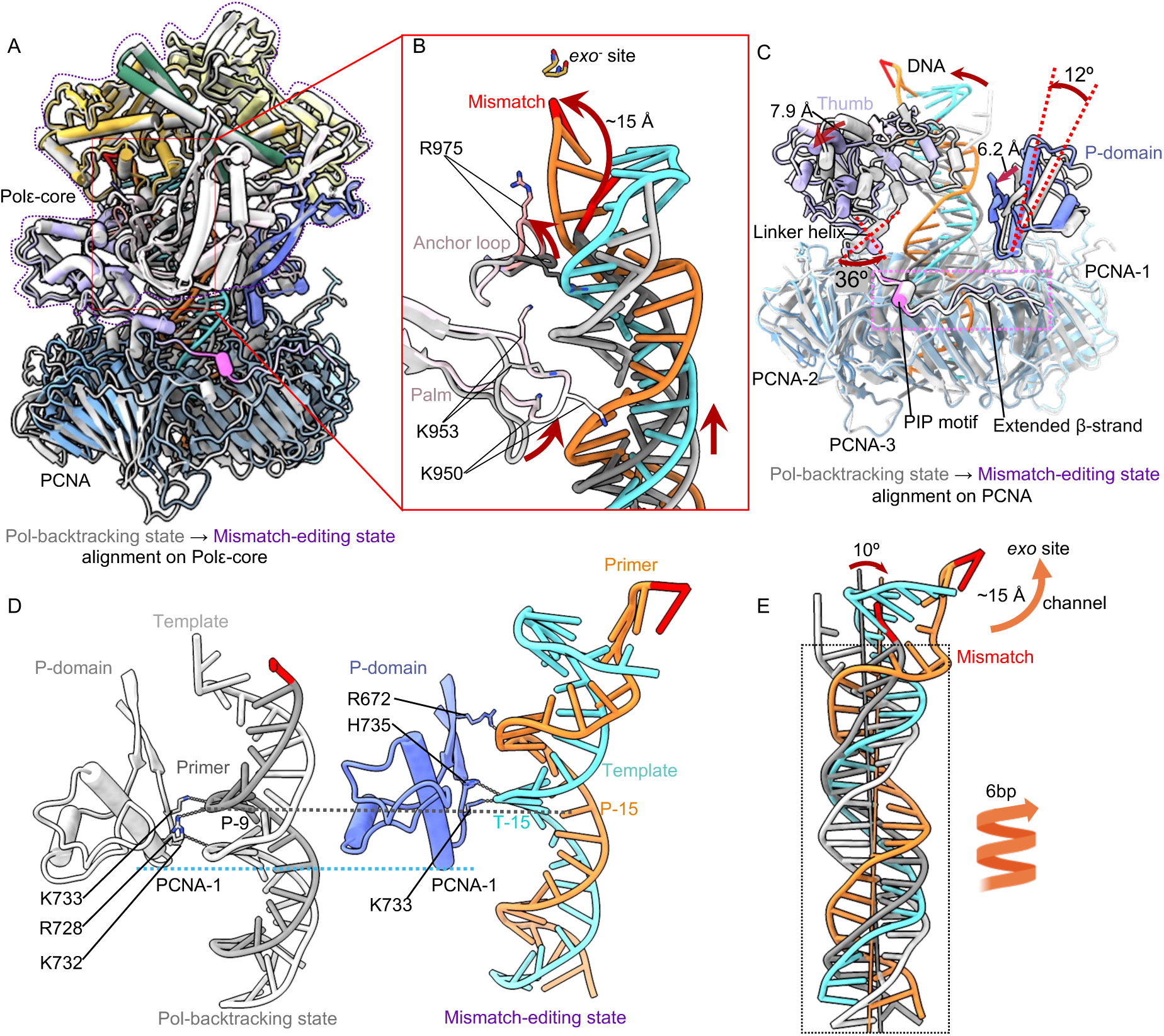
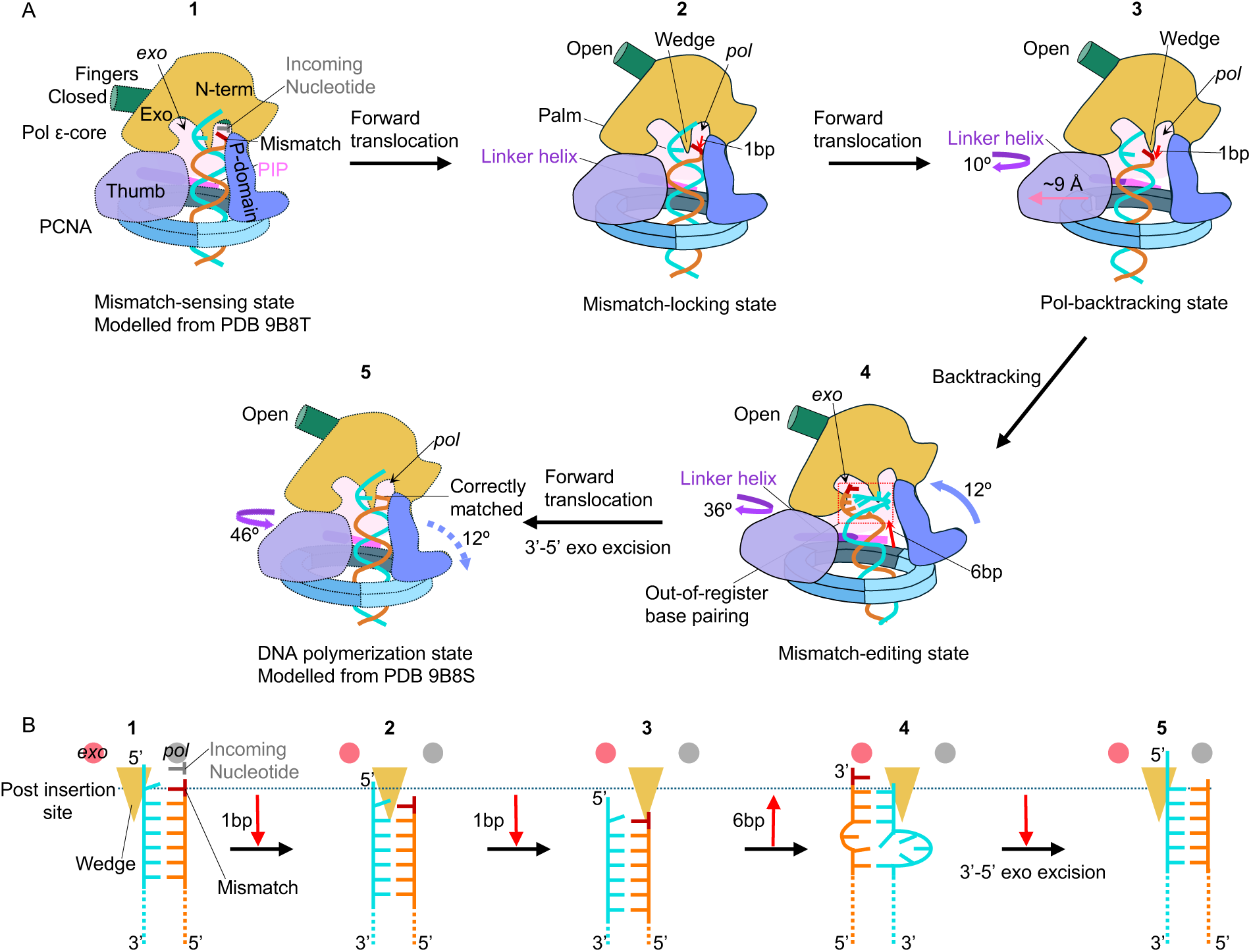
Conformational changes of Polε–PCNA induce the mismatched DNA translation during backtracking. (A) Comparison of the Pol-backtracking (gray) and mismatch-editing (color) states aligned on Polε-core. (B) Close-up view of the T/P and surrounding region. Red arrows indicate movements between the two states. Key residues are in sticks and labeled. (C) Structural changes near the Polε-PCNA interface. The top region of Polε-core is omitted for clarity. (D) Side-by-side comparison of the P-domain and T/P interaction in the two states. DNA-interacting residues are in sticks and labeled. (E) Comparison of the T/P in the two states reveal an overall translocation. The DNA tilts 10° toward the *exo* activity site. The duplex region spirals up by 6 bp, and 6 bp are unwound from the top primer 3′-end, enabling the primer to move 15 Å through exo channel into the *exo* site.

The interfaces between the Polε P-domain and PCNA-1 and between the Polε PIP motif plus the extended β-strand with PCNA-3 are maintained during the transition. But the thumb domain shifts laterally by 7.9 Å above PCNA-2. This large shift is driven by a ∼36° rotation of the linker helix connecting the thumb and the PIP motif. The P-domain tilts 12° toward DNA by pivoting on the bottom region of Polε that contacts PCNA, leading to a 6.2 Å shift of the P-domain β-strands such that they interact with the DNA minor groove (**Fig. 5C**). Thus, the bottom region of the Polε P-domain binds DNA in the Pol-backtracking state but it is the upper region of the P domain β-strands that bind DNA in the mismatch-editing state (**Fig. 5D**). Importantly, this moves the DNA up along with the P-domain during the transition correlating with the observed extensive unwinding. Correspondingly, Lys-733 interacts with the P-9 base in the Pol-backtracking state and with the T-15 base in the mismatch-editing state (**Fig. 5D**), consistent with the DNA spiraling up by 6 bp during the transition as well as 6-bp melting and rearrangement in the editing state (**Fig. 5E**). During the transition, DNA tilts by 10° and the Polε–PCNA holoenzyme backtracks, enabling the mismatched primer 3′-end to move by 15 Å to reach the *exo* site (**Fig. 5E**). These actions result in twice the unwinding observed in Pol editing modes in the absence of connection to processivity clamps.

## Discussion

Recent studies on the polymerizing state of Polε holoenzyme have shown that Polε-core engages with PCNA through a three-point interface, interacting with all three protomers of the PCNA clamp(24–26). However, it has been unclear if PCNA remains fully engaged during Polε proofreading, and if so, how the sliding clamp constrains the T/P movement while allowing the primer 3′-end to travel the long-distance from the *pol* to the *exo* site. PCNA is more than a processivity factor and is an integral part of a replicative Pol holoenzyme. Therefore, PCNA is expected to play an important role in the proofreading mechanism. But in the recent cryo-EM analysis of Polε proofreading, PCNA was largely dissociated from Polε in the presence of a pre-existing mismatched T/P with a 23-bp dsDNA region (25). The dissociation could be part of the proofreading mechanism to confer more freedom to Polε relative to the T/P as they negotiate the *pol*-to-*exo* transition, but Polε dissociation from PCNA might have been caused by the short duplex region of the T/P substrate used in that study. In the current study we used a longer duplex region and observed a 6-bp unwinding during Polε backtracking, and Polε remains attached to PCNA. Indeed, the shorter duplex region used in the previous study would not have been able to observe this consistent Polε–PCNA contact. Considering that the major interactions between Polε-core and PCNA observed in the polymerizing state are in fact retained throughout the proofreading process when using the longer duplex in this study, it seems likely that the proofreading process occurs as described in this report, although it is possible that there exist multiple proofreading pathways.

### *Bona fide* proofreading requires both PCNA and *in-situ*-generated mismatch

Remarkably, we found that the tightly coupled PCNA imposes such strong constraints on Polε that a T/P with a pre-existing mismatch could not insert from outside of the enzyme into the exo site. This observation calls into question the previous structural studies on the proofreading mechanism of other DNA polymerases that are known to function with a DNA sliding clamp (39–44).

We succeeded in capturing the proofreading states in the context of the holoenzyme only when the mismatch was synthesized in situ in the *pol* site of Polε. To our knowledge, this is the first time that DNA polymerase proofreading intermediates have been visualized in a near native environment in which the polymerase is complexed with the sliding clamp and the mismatch is first produced inside the *pol* site and then travels to the *exo* site. Under these more physiologically relevant conditions, we found that the human Polε proofreading mechanism is strikingly different from previous observations using preformed mismatched DNA that does not require Pol–clamp complex.

### Proposed proofreading process by the human Polε–PCNA holoenzyme

Morphing the three proofreading intermediates observed here provides a plausible view of the *pol*-to-*exo* translocation process of a mismatched T/P DNA that is made in the Polε active site and is then transferred to the exo site for removal of the mismatch (**Movie S1**). We therefore propose the following proofreading mechanism for the human Polε (**Fig. 6**). It is known that proofreading is initiated (step 1) when a mismatched base is synthesized at the *pol* site and fails the Watson-Crick fidelity check, and there have been studies on how this fidelity check might work (52–54). But for the earliest state captured in this report Polε has shifted forward by 1 bp on the template and thus has passed the checkpoint step, and is therefore at step 2, the mismatch-locking state (**Fig. 6**). Due to the T/P having backtracked from the *pol* site by 1 bp, the fingers domain is now switched to the open conformation (Pol inactive). This 1-bp-mismatch displacement is important for preventing potential Pol extension on the mismatch. In step 3, Polε rotates slightly above the PCNA, accompanied by a 10° outward rotation of the linker helix between the PIP motif and the thumb domain, and a 20° tilt of the DNA to shift the thumb domain outwards by 9 Å, making room for DNA translocation (**Fig. 3B**). These changes result in the anchor loop that appears involved in separating the primer 3’-end from template (51). In step 4, the Polε P-domain tilts by 12° and the linker helix rotates by 36° to drive T/P unwinding by 6 bp and move the mismatched primer 3′-end by 15 Å towards the *exo* site (**Fig. 5C-E**). The 6-bp T/P melting is the longest unwinding for proofreading observed thus far, as previous studies observed melting of only 3 bp during proofreading in Polε (25) and other B-family DNA polymerases (39–44). In the last step, step 5 which is not observed in this study, the mismatched base is expected to be cleaved by the 3′-5′ exo activity before the primer returns to the *pol* site to resume DNA synthesis (**Fig. 6**).

**Fig. 6.**
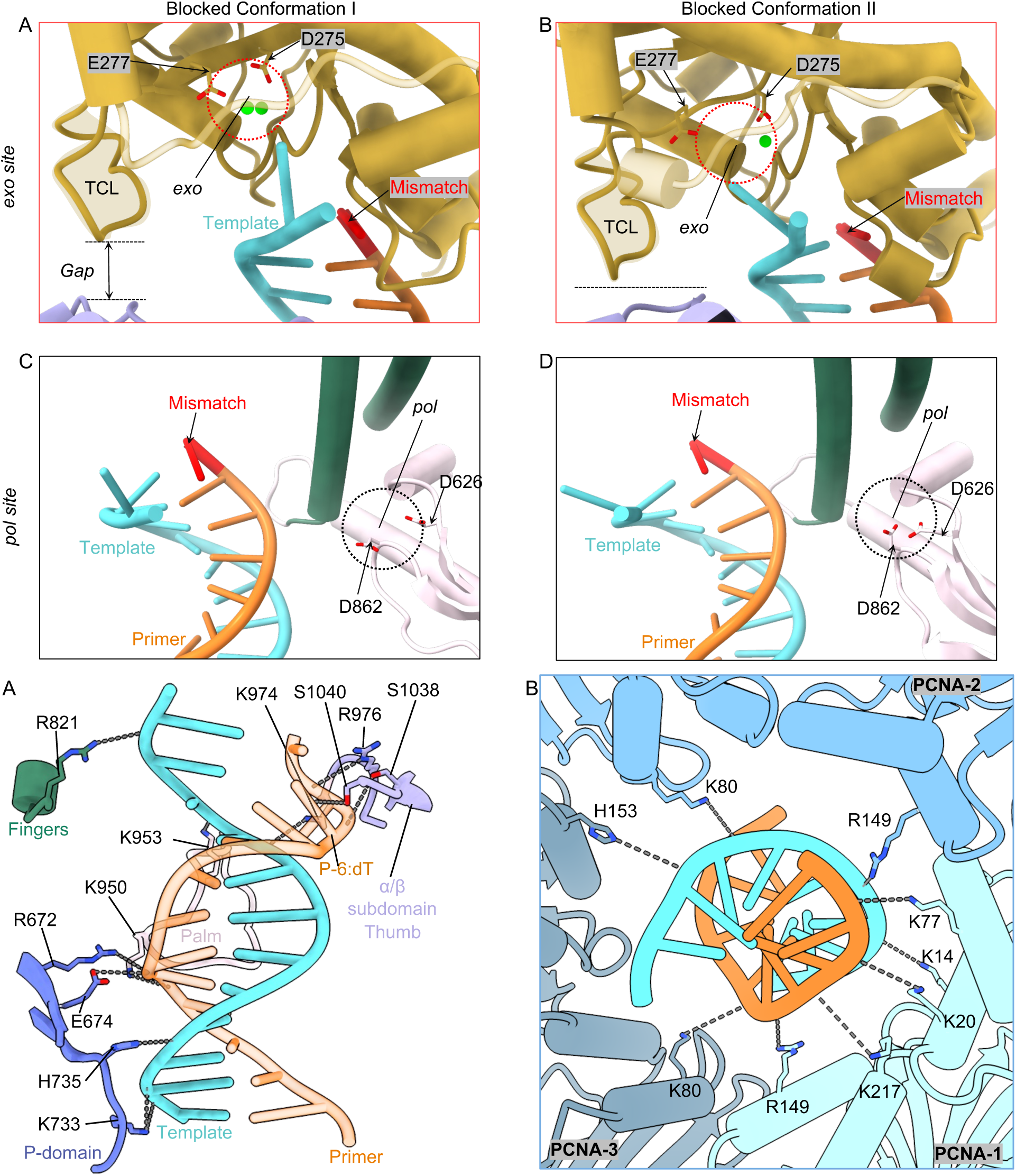
Model of the Polε–PCNA holoenzyme proofreading process. (A) Schematic representation of all five steps that are built based on experimental structures of the human Polε– PCNA holoenzyme. In step 1, Polε incorporates a mismatched base in the *pol* site and senses the mismatch via a Watson-Crick base-pairing checkpoint. This is modeled by Polε structure in the Pol state (PDB ID 9B8T). In step 2, Fingers domain flips up to an open position to arrest the pol activity, and the holoenzyme moves away from the mismatched 3′-end by 1 bp to prevent additional base incorporation. In step 3, the Polε linker helix rotates 10° and the thumb domain moves outwards by 9 Å. The movements put pressure on the 1-bp backtracked T/P. In step 4, the P-domain tilts 12° against the PCNA, and the linker helix rotates 36°, causing the holoenzyme to backtrack by 6 bp and unwind 6 bp from the primer 3′-end. The 6 unwound bp rebind and form 4 out-of-register bp and to insert the mismatched primer 3′-end into the *exo* site. In step 5, the primer 3′ mismatch is excised by the exo activity. Next (not shown), in an expected reverse process not yet well characterized, the holoenzyme returns the T/P to the *pol* site and resumes DNA synthesis. (B) Schematic describing the mismatch DNA translocation during the Polε–PCNA holoenzyme proofreading process. The horizontal dashed line indicates the post insertion site of Polε. The movement of DNA between each step is indicated by red arrows.

### Implication of the P-domain involvement in Polε proofreading

The eukaryotic Polε has undergone two remarkable inventions during evolution: formation of the POLE1 catalytic domain fused to another B family noncatalytic POLE1 CTD that tethers POLE to the replicative helicase, and addition of the P-domain in the catalytic NTD. Recent studies have shown that the P-domain enhances Polε processivity by directly binding to DNA and PCNA (23–26), and the P-domain also stabilizes the intrinsically mobile Ctf18 AAA+ domain to help PCNA loading onto the leading strand DNA by the Ctf18-RFC clamp loader (34, 62–64). This study now reveals a third function of the P-domain in T/P melting and repositioning during mismatch proofreading (**Fig. 5C-D**). Because the P-domain is unique to Polε, this raises the question whether the proofreading mechanism we have described above is unique to Polε. Perhaps other replicative DNA polymerases may use a different proofreading mechanism, possibly by dissociating from their processivity clamp. Structural analysis of other polymerases in the presence of a sliding clamp and use of a mismatch generated in situ will be needed to clarify this question.

### Implications of out-of-register base-(mis)pairing of melted T/P

An important discovery of this study is the out-of-register base pairing or base mispairing of the 6 melted base pairs. In the previously reported Pol proofreading state, 3 base pairs are unwound, and the unwound regions of template and primer are kept separate by the long β-hairpin that protrudes downwards from the exo domain (55, 58). But the corresponding β-hairpin loop is uniquely short in Polε and does not reach the template to keep it from rebinding to the primer, and this out-of-register rebinding leaves only the single mismatched 3′-nucleotide in the *exo* site (**Fig. 4E-F**). This represents a pre-excision stage of the exo-editing model in which Polε is unable to continue cutting the correct 3′ primer bases. It is currently unclear if such T/P proofreading configuration is adopted in other Pols or whether this process is unique to Polε–PCNA. Regardless, this configuration may be related to the long-recognized but mysterious “replication slippage” phenomenon (65–67). Slippage frequently occurs in DNA regions containing homo-nucleotide runs – stretches of DNA with an identical nucleotide. We speculate that out-of-register rebinding of the melted primer and template regions during proofreading will result in perfect base pairing in a homo-nucleotide run, and such base pairing is harder to break up for realignment (in-register base pairing) when the exo-edited T/P returns to the Pol site. In these cases, failure of T/P realignment might then introduce replication slippage upon resumption of DNA synthesis in the *Pol* site.

This study represents the first revelation of a bona fide proofreading process of the human Polε-PCNA holoenzyme. Mutations in the Polε exo domain compromise replication fidelity and are drivers for tumorigenesis in a wide range of human cancers (68, 69). Our study hence highlights those mutations at the interface between Polε and PCNA may also affect its proofreading activity and could be involved in cancer progression. We also note that the approach taken here, using in situ mismatch formation, can be used to interrogate proofreading mechanisms of many other DNA polymerases.

## Materials and methods

Details on the materials and methods for the expression and purification of human Polε, Polε-core and PCNA are provided in the **SI Appendix**. As a control for the structural analysis of the in-situ formation of a mismatch, Polε or Polε-core and PCNA were directly mixed with DNA containing a pre-existing mismatch. For in situ formation of a mismatch, the PCNA and Polε were mixed with a flush primed DNA followed by adding dNTPs that result in a mismatch. Only the in-situ method provided the physiologically relevant proofreading intermediates. The detailed methods off assembly, grid formation, data collection, 3D map reconstruction, model building and refinement are provided in the **SI Appendix**.

### Data, Materials, and Software Availability

Cryo-EM structural data have been deposited in the Electron Microscopy Data Bank (https://www.ebi.ac.uk/pdbe/emdb/) and the Protein Data Bank (https://www.rcsb.org) under the following accession codes: Human Polε–PCNA–T49/P35 (pre-existing mismatch) in the blocked conformation I (EMD-49302 (70) and 9NE9 (71)), human Polε–PCNA–T49/P35 (pre-existing mismatch) in the blocked conformation II (EMD-49303 (72) and 9NEA (73)), human Polε-core– PCNA–T47/P29 (with in situ generated mismatch) in the Mismatch-locking state (EMD-49301(74) and 9NE8 (75)), human Polε-core–PCNA–T47/P29 (with in situ generated mismatch) in the Pol-backtracking state (EMD-49300 (76) and 9NE7 (77)), and human Polε-core–PCNA–T47/P29 (with in situ generated mismatch) in the Mismatch-editing state (EMD-49299 (78) and 9NE6 (79)).

## Acknowledgments

Cryo-EM data were collected at the David Van Andel Advanced Cryo-Electron Microscopy Suite of Van Andel Institute. We thank G. Zhao and X. Meng for their assistance with data collection. This study was supported by the US NIH grants GM148159 (M.E.O) and GM131754 (H.L.), Howard Hughes Medical Institute (M.E.O), and Van Andel Institute (H.L.).

## Author contributions

F.W., Q.H., M.E.O., and H.L. designed the research; F.W. and Q.H. performed experiments; F.W. contributed new reagents/analytic tools; Q.H., F.W., and H.L. analyzed data; and F.W., Q.H., M.E.O., and H.L. wrote the manuscript.

---

**SI Appendix, Table S1.**
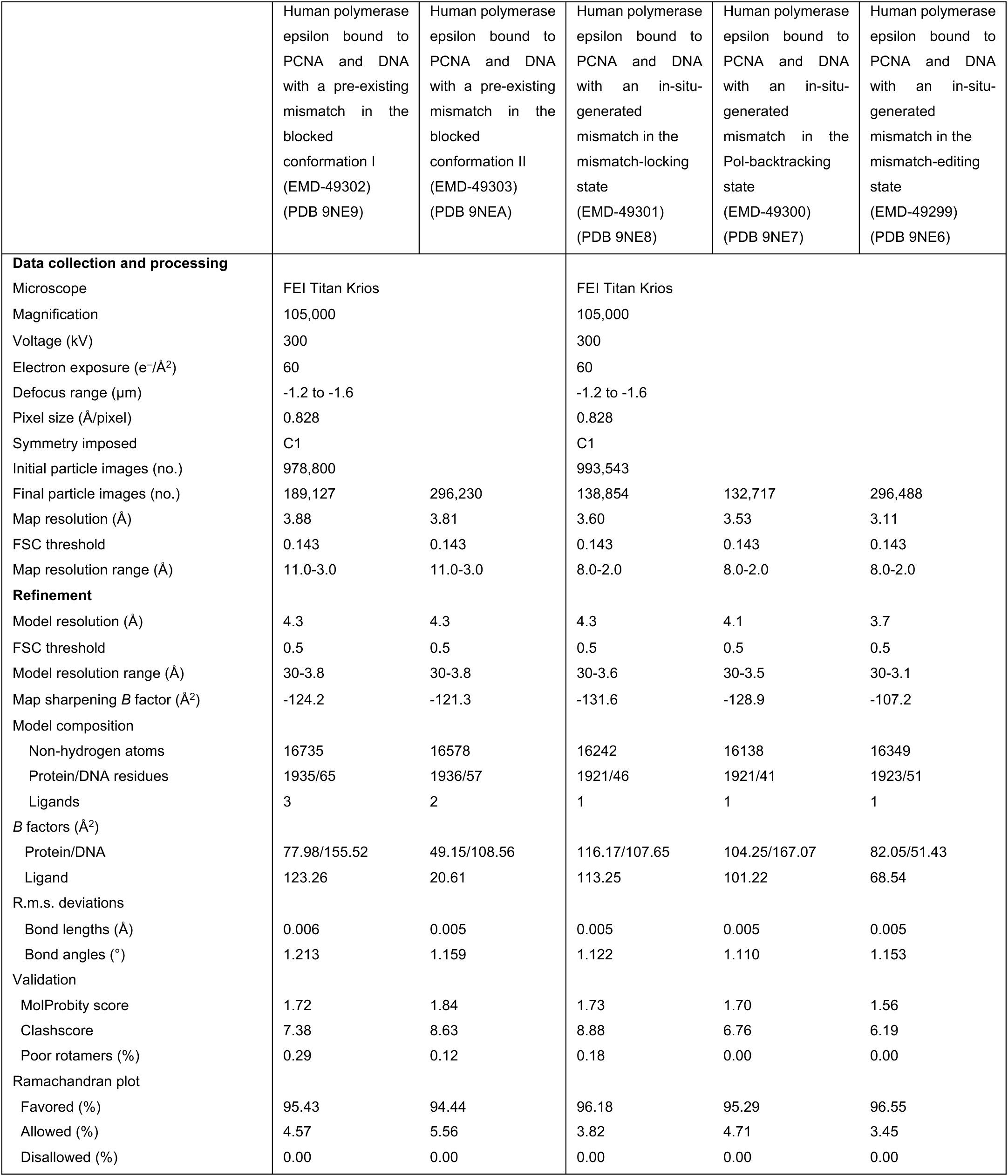
Cryo-EM data collection, refinement, and validation statistics.

**SI Appendix, Fig. S1.**
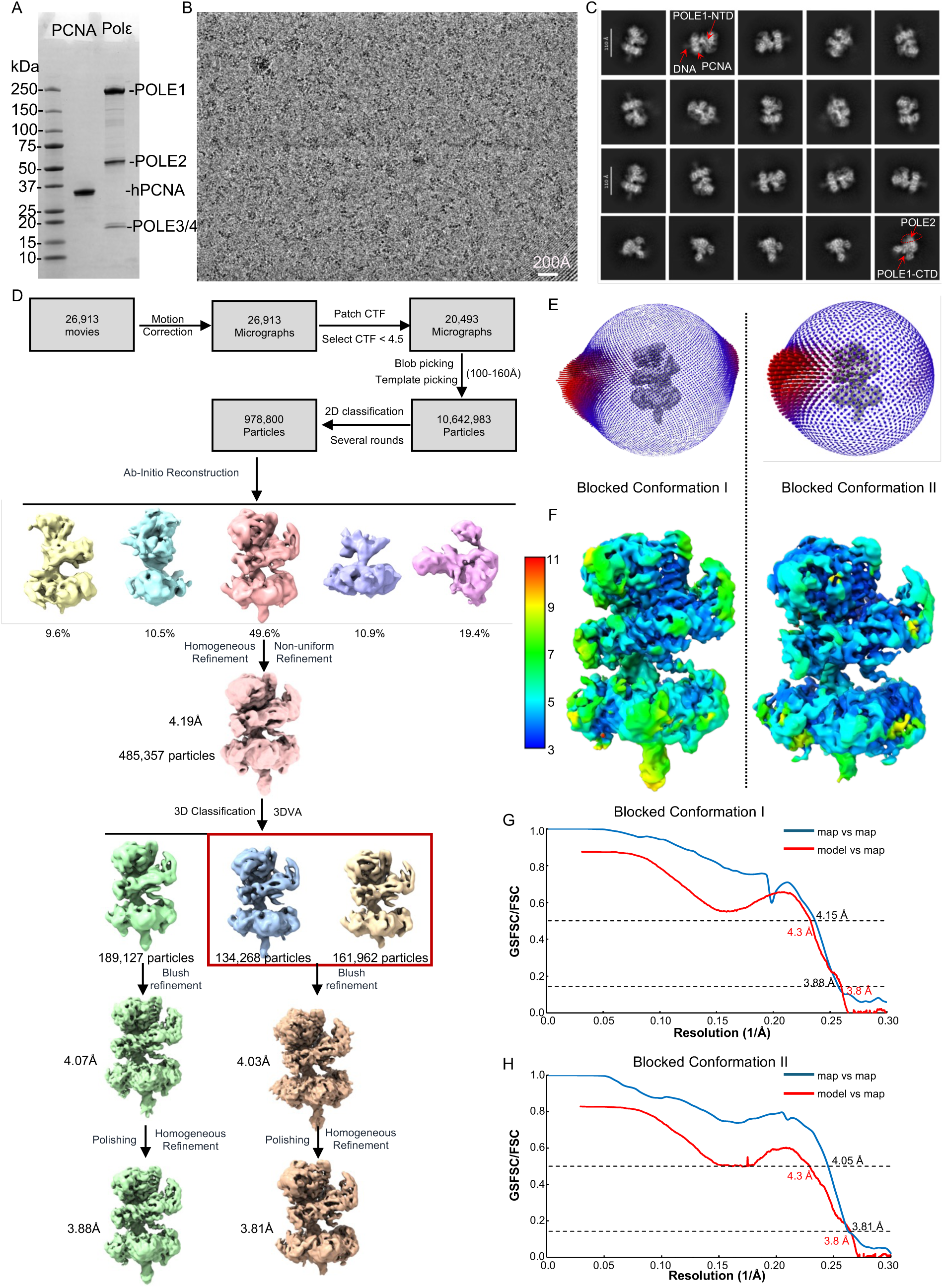
Cryo-EM data processing of the Polε–PCNA–T49/P35 ternary complex (with pre-existing mismatch). (A) SDS-PAGE gel of purified wild-type human Polε and PCNA. (B) Typical raw micrograph of the in vitro assembled complex. A total of 26,913 micrographs were recorded. (C) Selected 2D class averages in various views. (D) Workflow of cryo-EM data processing and 3D reconstruction in CryoSPARC (version 4.5.1), leading to the two 3D EM maps at 4.01 Å and 3.95 Å, respectively. (E) Angular distribution of particle images contributing to the two final 3D reconstructions. (F) Color-coded local resolution maps of the ternary complex in two blocked conformations. (G-H) Gold standard Fourier shell correlation (GSFSC) of the two EM maps (blue) and the model-to-EM map correlation curve (red) of the ternary complex in two blocked conformations.

**SI Appendix, Fig. S2.**
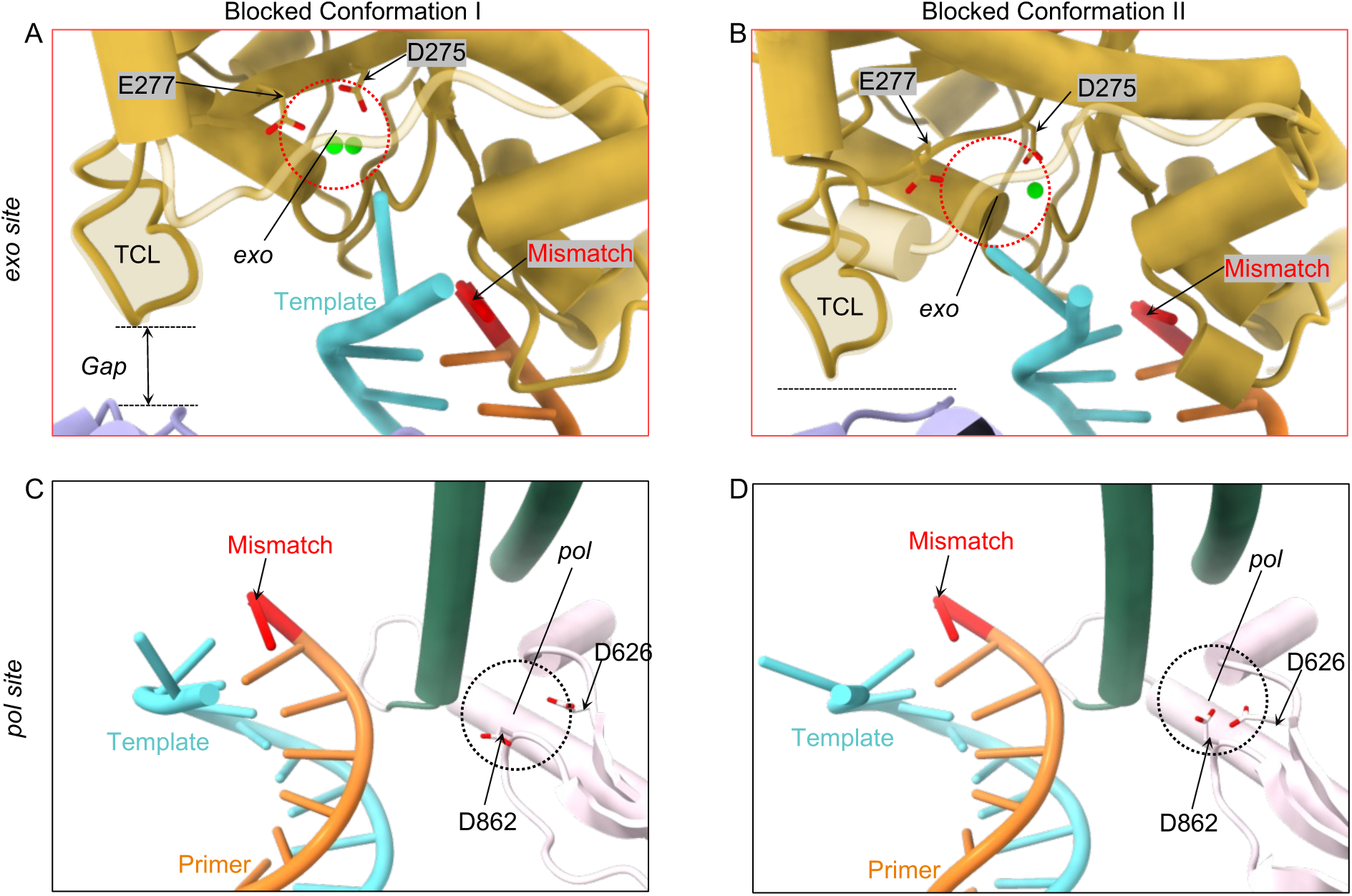
DNA positions in the Polε–PCNA–T49/P35 (with pre-existing mismatch) in the two blocked conformations. (A-B) The mismatched primer 3′-end (red stick) with respect to the Polε *exo* site (dashed red circle) of the holoenzyme in blocked conformations I (A) and II (B). The mismatched primer 3′-end is far away from the *exo* site. There is a gap between the exo and thumb domains in the blocked conformation I, but the gap is closed via a TCL loop in the blocked conformation II. The gap may facilitate the template strand to enter a channel leading up to the *exo* site. (C-D) The mismatched 3′-end (red stick) with respect to the *pol* site (dashed black circle) in blocked conformation I (C) and conformation II (D). The pol site contains no catalytic metal ions to coordinate the catalytic residues D626 and D862 residues, and the mismatched 3′-end is far from the *pol* site.

**SI Appendix, Fig. S3.**
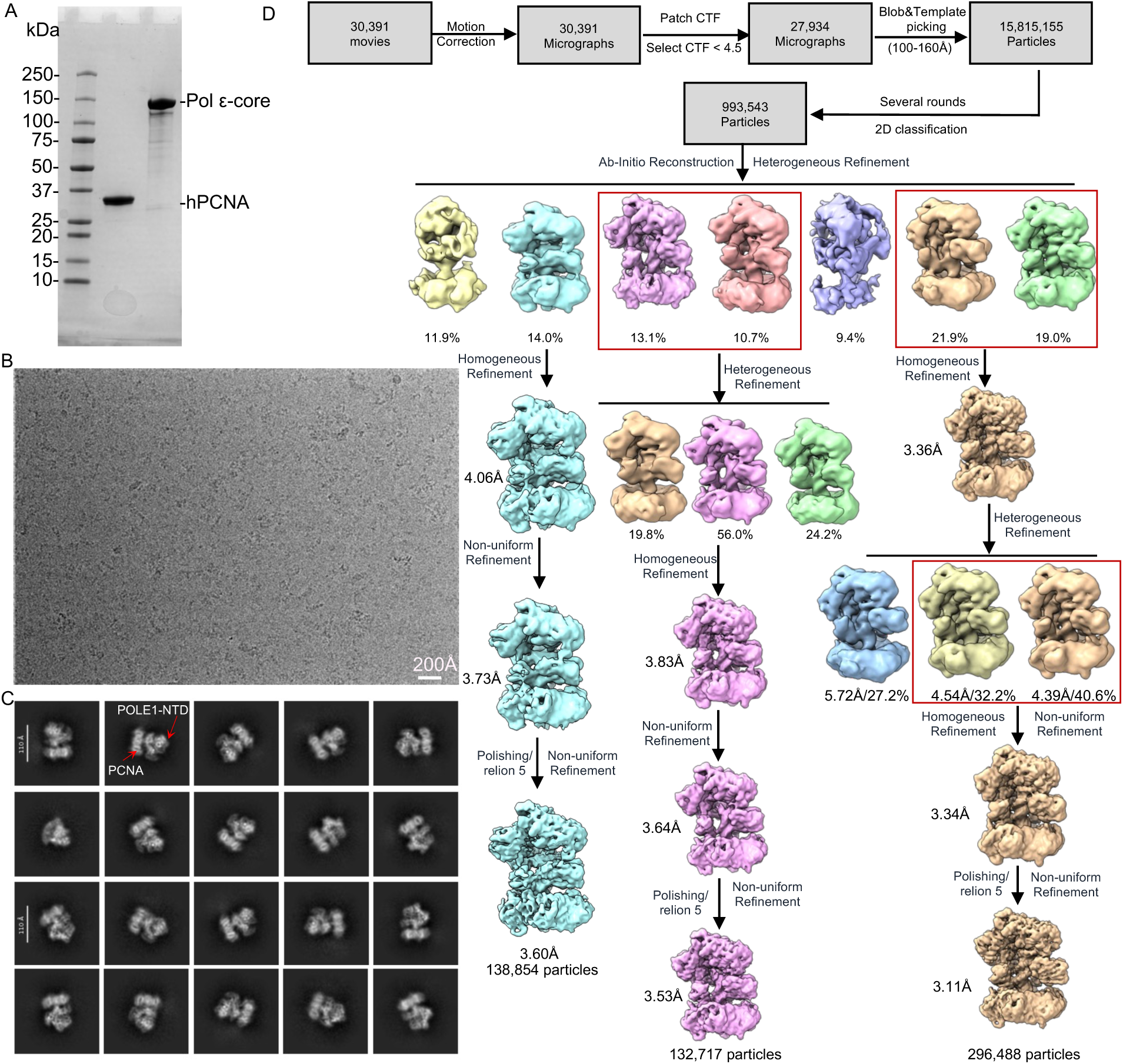
Cryo-EM data processing of the Polε–PCNA–T47/P29 (with in-situ-generated mismatch). (A) SDS-PAGE gel of purified human Polε-core exo^-^ and PCNA. (B) Typical raw micrograph out of the total 30,391 recorded movies. (C) Selected 2D class averages in various views. (D) Workflow of cryo-EM data processing and 3D reconstruction in CryoSPARC (version 4.5.1), leading to three 3D EM maps at 3.60 Å, 3.53 Å, and 3.11 Å average resolution, respectively. The final Bayesian polishing was performed by Relion-5.

**SI Appendix, Fig. S4.**
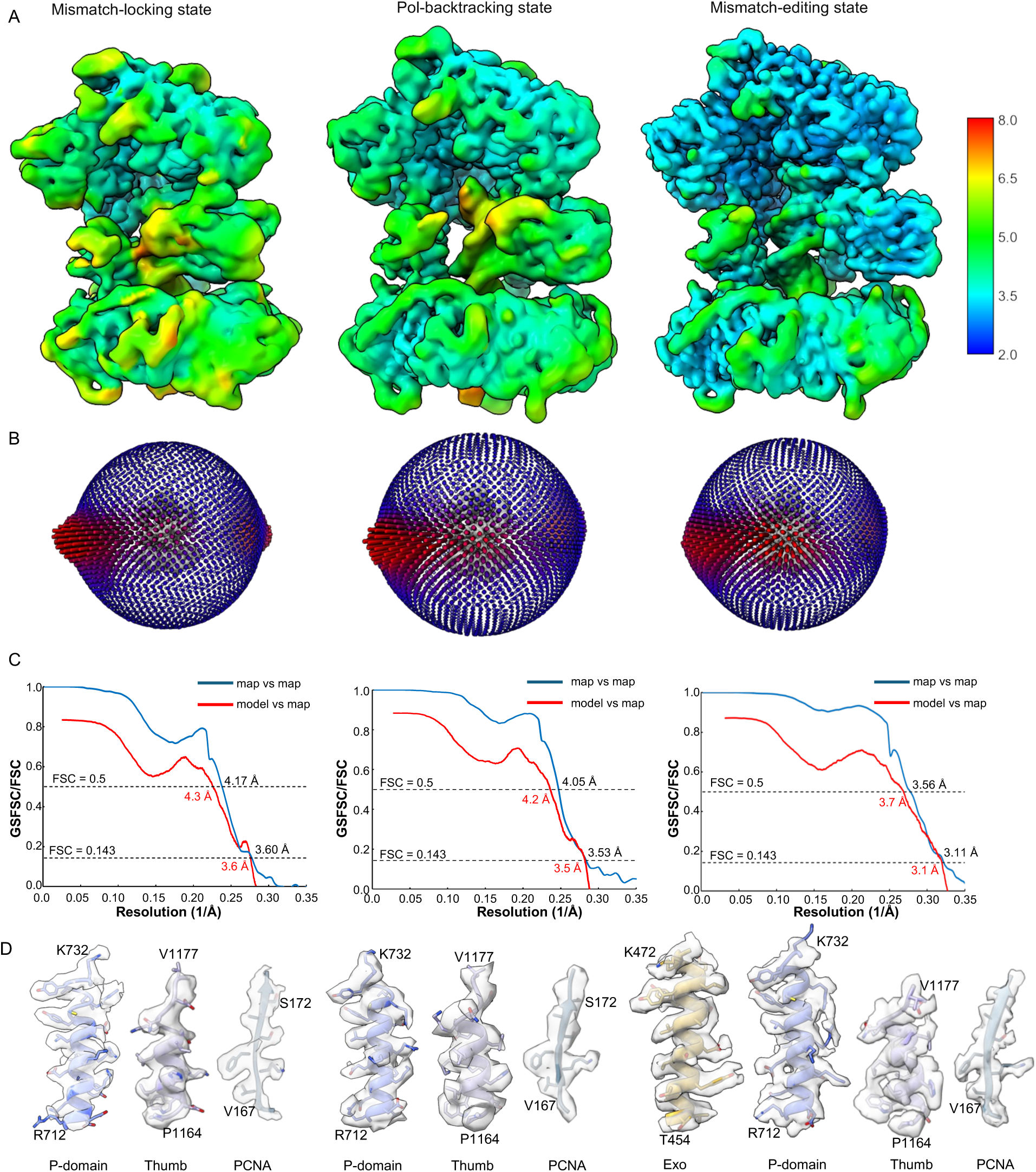
Resolution estimation of the 3D EM maps of Polε–PCNA–T47/P29 (with in-situ-generated mismatch) in three proofreading states. (A) Color-coded local resolution maps of three proofreading states. (B) Angular distribution of particle images used in the final 3D reconstructions. (C) Gold standard Fourier shell correlation (GSFSC, blue) and the map-to-model correction (red) curves. (D) EM densities in three selected regions of each of the three proofreading states.

**SI Appendix, Fig. S5.**
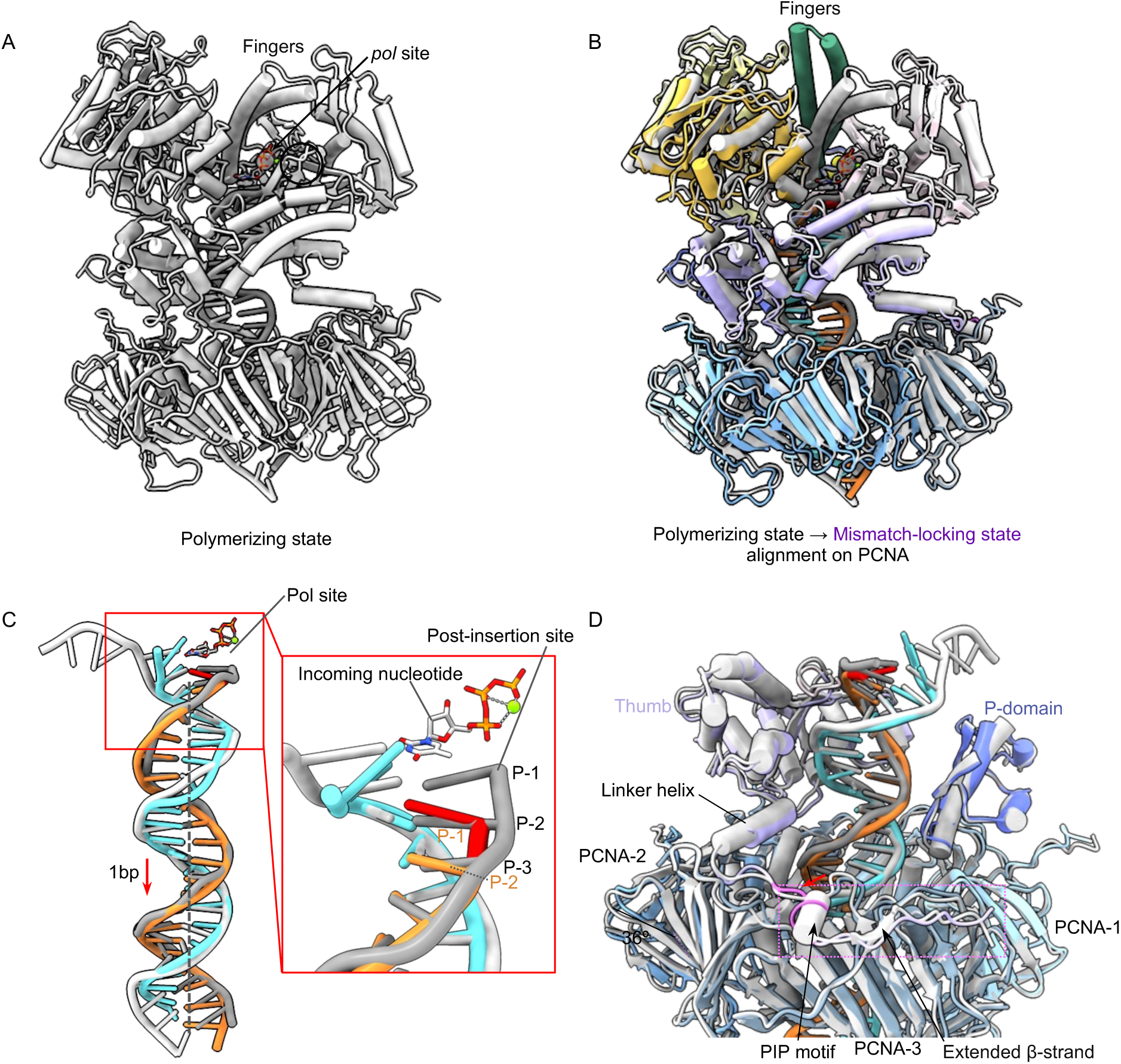
Conformational changes of the human Polε holoenzyme from the polymerization to the mismatch-locking state. (A) Structure of the reported polymerization state (PDB ID 9B8T) of Polε–PCNA with a matched T/P. (B) Superimposition of the polymerization (gray) and mismatch-locking (color) states reveals that the Fingers domain transitions from the closed (gray) to open (dark green) conformation. (C) Superposition of the T/P reveals a 1-bp downward shift from the polymerization to the mismatch-locking state. (D) No major changes occur at the Polε-PCNA interface region from the polymerization (gray) to the mismatch-locking (color) states.

**SI Appendix, Fig. S6.**
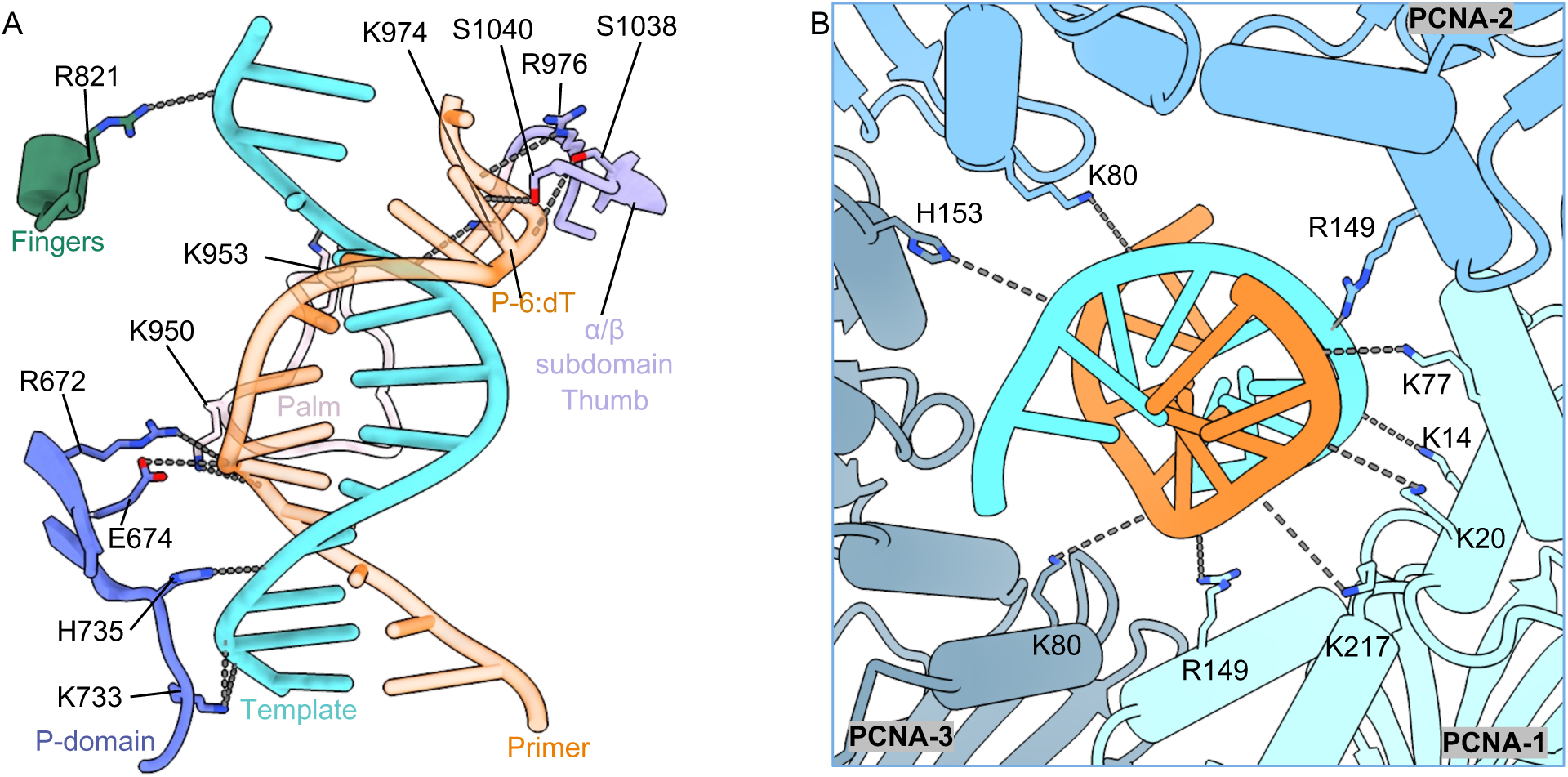
The duplex region of the T/P is well stabilized by the Polε–PCNA holoenzyme during mismatch-editing. (A) The T/P duplex region above PCNA interacts with residues from the Fingers, P-domain, and thumb domain. (B) The T/P duplex region inside PCNA is slightly tilted and interacts with 9 positively charged residues.

## Supplementary Information

## Materials and Methods

### Molecular cloning and DNA oligomers

The cDNAs of human Polε (POLE1-4) were acquired from DNASU Plasmid Repository, and the cDNA of human PCNA was obtained from Addgene. The cDNA of POLE1(Gene ID: 5426) was tagged with an 6xHis sequence at N-terminus and inserted into the pFL multi-gene expression vector(1). And the cDNAs of POLE2 (Gene ID: 5427, with an N-terminal 3xFLAG tag), POLE3 (Gene ID: 54107), and POLE4 (Gene ID: 56655, with an N-terminal 3xFLAG tag) were cloned into a second pFL multi-gene expression vector(1). The truncated POLE1 (1–1200 aa, Polε-core), containing a tandem N-terminal 3×FLAG-6×His tag, was cloned into the pFastBac vector. The cDNA of human PCNA (Gene ID: 5111) with an N-terminal 6xHis tag was cloned into the pET28a vector. The exo^-^ site mutations (D275A/E277A) of POLE1 or Polε-core were generated by PCR-based mutagenesis. All constructs were sequenced to ensure that mutations were not introduced during PCR and cloning.

The DNA substrates used in the cryo-EM structural studies were chemically synthesized by Eurofins Genomics. The correctly matched T/P DNA substrate (T47/P29) includes a 29-nt primer strand (5′-TGAGGTTCAGCAAGGTGATGCTTTAGATT-3′) and a 47-nt template strand (5′-GCCAGCAG CAAAGT<u>G</u>AAAAATCTAAAGCATCACCTTGCTGAACCTCA-3′). The mismatched T/P DNA (T49/P35) contains a 35-nt primer strand (5′-TGAGGTTCAGCAAGGTGATGCTTTAGATTTTTCA*<u>T</u>-3′) and a 49-nt template strand (5′-GAGCCAGCAGCAAAGTGAAAAATCTAAAGCATCACCTTGCTGAACCTCA-3′).

A phosphorothioate bond (*) was also introduced between the last two nucleotides of the 35-nt primer to prevent cleavage of the mismatched primer 3′-end. For controls, a standard 35-nt primer strand without the phosphorothioate modification was also prepared. But the T49/P35 substrate with or without the modification led to the same Polε structure. Primer and template oligonucleotides were mixed in equimolar amounts to achieve a final concentration of 100 μM in an annealing buffer (20 mM HEPES, pH 7.5, 50 mM NaCl, and 0.5 mM EDTA). The mixture was heat denatured at 95°C for 10 minutes and then gradually cooled to room temperature. Finally, the annealed P/T DNA were stored at –20°C until use.

### Protein expression and purification

Wild-type Polε or Polε exo^-^, featuring a 6xHis tag at the N-terminus of POLE1 and 3xFLAG tags at the N-termini of POLE2 and POLE4, was expressed using the Bac-to-Bac Baculovirus expression system (Thermo Fisher), following a protocol similar to that used for the EXO mutant of Polε (2). Sf9 cells were co-infected with baculoviruses encoding POLE1 and the POLE2-4 subcomplex and incubated at 27°C with shaking at 115 rpm for 65-72 hours. The harvested insect cells were sonicated in lysis buffer (25 mM HEPES, pH 7.5, 250 mM NaCl, 1 mM MgAc, 5% glycerol, and one tablet of EDTA-free protease inhibitor cocktail). The lysate was centrifuged at 125,440 × g for 1 hour using a Ti-45 rotor, and the supernatant was incubated with 0.8 mL of FLAG antibody-conjugated beads at 4°C for 2-3 hours. After washing the beads twice with 50 mL lysis buffer, the bound proteins were eluted with 8 mL lysis buffer supplemented with 0.2 mg/mL FLAG peptide. The eluted protein was concentrated using an Amicon centrifugal concentrator (100 kDa cutoff) and further purified by size-exclusion chromatography (Superose 6 Increase, GE Healthcare) in buffer containing 25 mM HEPES, pH 7.5, 200 mM NaCl, 1 mM MgAc, and 1 mM DTT. The purified Polε was concentrated to 3.7 mg/mL and stored at –80°C. Polε-core or Polε exo^-^ was expressed and purified following the same protocol used for full-length Polε. Human PCNA was expressed in *E. coli* BL21 as previously described(3). Briefly, PCNA expression in *E. coli* was induced by 0.3 mM IPTG at 16°C for 18 hours. Cells were harvested and resuspended in lysis buffer (25 mM HEPES, pH 7.5, 200 mM NaCl, 5% Glycerol). Cell lysates were obtained by homogenization using an SPX Corporation homogenizer and subsequently clarified by centrifugation at 34,572 x g for 1 hour at 4°C. His-tagged PCNA was purified using a Ni-NTA column (Cytiva), followed by size-exclusion chromatography on a Superdex 200 column (GE Healthcare) in buffer containing 25 mM HEPES (pH 7.5), 200 mM NaCl, and 1 mM DTT. Purified PCNA was concentrated to 3.0 mg/ml and stored at –80°C.

### Cryo-EM sample preparation and data collection

To assemble the *in vitro* ternary complex of human Polε–PCNA–DNA (pre-existing mismatched T/P), we first mixed 1 µM Polε/Polε-core, 3 µM PCNA, 1.1 µM mismatched DNA (T49/P35), and 0.5 mM dTTP at room temperature for 10 minutes and incubated the mixture on ice for 2 hours prior to grid preparation. The simultaneous addition of all three components only produced a ternary complex in the blocked state. We next changed the assembly scheme by first mixing PCNA and DNA, then adding Polε or Polε-core and dNTPs. But this produced the same blocked states. To address the preferred particle orientation issue, we added 0.02% octyl β-D-glucoside (β-OG) into the sample solution immediately before grid vitrification. We did not perform the third assembly scheme of first mixing Polε or Polε-core with mismatched DNA, followed by addition PCNA. Because this mixing scheme was used previously(4) which resulted in Polε/Polε-core interacting with the mismatched DNA prior to PCNA, a scenario we wanted to avoid.

To assemble the ternary complex of human Polε–PCNA with an in-situ-generated mismatched T/P DNA, we mixed 1 µM Polε exo^-^/Polε-core exo^-^, 3 µM PCNA, and 1.1 µM correctly-matched DNA (T47/P29) and incubated the mixture on ice for 2 hours, then added 0.5 mM dTTP and incubated the reaction mixture at room temperature for 3 minutes to allow Polε to extend the primer by four bases and generate a terminal G•T mismatched base pair in situ in the *pol* site. For cryo-EM grid preparation, holey carbon grids (Quantifoil Au R1.2/1.3, 300 gold mesh) were glow-discharged in an Ar/O_2_ mixture for 30 seconds using a Gatan 950 Solarus plasma cleaner. A 3 µL aliquot of each final reaction mixture containing 0.02% β-OG was applied to the freshly treated grids. Vitrification was performed using a Vitrobot Mark IV system (Thermo Fisher Scientific) with the following settings: blot time of 3 s, blot force of 3, wait time of 5 s, sample chamber temperature of 6 °C, and chamber relative humidity of 100%. Grids were then flash-frozen in liquid ethane cooled by liquid nitrogen.

Cryo-EM data were collected using a 300 kV Titan Krios microscope, operated via SerialEM in multi-hole mode(5). Images were all acquired at 105,000× magnification, with defocus values ranging from –1.2 to –1.6 µm. The data were captured on a Gatan K3 direct electron detector in super-resolution mode, with a pixel size equivalent to 0.414 Å at the specimen level. Each exposure lasted 1.0 second, during which 50 frames were captured, resulting in a total dose of 60 e^−^/Å².

### Image Processing and 3D Reconstruction

For the ternary complex of Polε–PCNA-T49/P35 (pre-existing mismatch), a total of 26,913 raw movie micrographs were collected and motion-corrected using MotionCorr 2.0(6) with 2× binning, yielding a pixel size of 0.828 Å/pixel. The motion-corrected micrographs were imported into cryoSPARC (version 4.5.1) for patch-based contrast transfer function (CTF) estimation(7), and 20,493 micrographs with CTF signals extending to 4.5 Å were selected for further processing (**Supplementary Fig. 1d**). Blob-based auto-picking (particle diameters of 100–160 Å) was implemented in cryoSPARC(7) to select initial particle images, which were used to generate a set of 2D classes for the next template-based particle picking. A total of 10,642,983 raw particles were automatically picked and 4× binned. After several rounds of 2D classification, particles with clear structural features were selected. In total, 978,800 selected particles were extracted with a box size of 320 pixels and used to compute five initial 3D models. Three low-quality 3D reconstructions were discarded as junks. One 3D model represented the noncatalytic form of Polε but failed to reach a good resolution upon further refinement. The remaining 3D model had good structural features and was selected for homogeneous and non-uniform 3D refinements, resulting in a 3D map that contained high conformational heterogeneity. Further 3DVA analysis (8) and 3D classifications were performed to generate three subclasses. One subclass was subjected to the Blush regularization refinement(9) to produce an intermediate 3D map at an overall resolution of 4.07 Å. The particles belong to this subclass were then imported into Relion 5 for Bayesian polishing Relion 5 (10), followed by non-uniform refinement in cryoSPARC, leading to a slightly improved final resolution of 3.88 Å. The remaining two subclasses were combined and subjected to Blush regularization refinement, generating an intermediate 3D map at 4.03 Å resolution. Further Bayesian polishing and non-uniform refinement in cryoSPARC improved the resolution to 3.81 Å.

For the ternary complex of Polε–PCNA–T47/P29 with an in situ-generated mismatch, a total of 30,391 raw movie micrographs were collected and motion-corrected using MotionCorr 2.0(6) with 2× binning and an effective pixel size of 0.828 Å/pixel. The motion-corrected micrographs were imported into cryoSPARC (version 4.5.1) for patch-based contrast transfer function (CTF) estimation(7), and 27,934 micrographs with CTF signals extending to 4.5 Å were selected for blob-based automatic particle picking (with particle diameters of 100–160 Å). The blob-picked particles were subjected to 2D classification resulting in a set of 2D averages with structural details expected of the Polε–PCNA particles. These 2D averages were used as templates for the next round of template-based particle picking. A total of 15,815,155 raw particles were automatically picked and 4× binned. After several rounds of 2D classification, particles with clear structural features of the Polε–PCNA complex were selected. In total, 993,543 selected particles were extracted with a box size of 320 pixels and used to compute seven initial 3D models, and selected 3D models with promising structural features were used as 3D template models for heterogenous 3D refinements against all particles. This resulted in seven 3D subclasses. Two low-quality 3D subclasses reconstructions were discarded as junk. One 3D subclass with good structural features for both Polε and PCNA was selected for homogeneous and non-uniform 3D refinements, resulting in a 3D map at 3.73 Å resolution. The 138,854 particles associated with this 3D map were imported into Relion 5 for Bayesian polishing (10), followed by non-uniform refinement in cryoSPARC, leading to an improved final resolution of 3.60 Å. Two 2D subclasses with the frayed DNA features were combined and subjected to another round of heterogenous refinement. This resulted in a major 3D class with good structural features, which was selected for homogeneous and non-uniform 3D refinements, resulting in a 3D map at 3.64 Å. The 132,717 particles contributing to this map were Bayesian polished in Relion 5 and homogeneous refined in cryoSPARC to produce the final 3.53 Å cryo-EM map. Two remaining 3D subclasses with similar structural features were combined and subjected to a homogenous refinement, resulting in a 3D map at an overall resolution of 3.36 Å. The 3D map has clear density in the Polε region but weaker density in the PCNA region. To address this, a new round of heterogeneous refinement was performed. By discarding a low-resolution 3D subclass and combining the two remaining 3D subclasses for further homogeneous and non-uniform 3D refinements, an improved 3D map at 3.34 Å was obtained. This map was subjected to Bayesian polishing in Relion 5 and non-uniform refinement in cryoSPARC, resulting in the map at 3.11 Å resolution.

### Model building, Refinement, and Validation

The cryo-EM structure of human Polε-core–PCNA with an open finger conformation of POLE1-NTD (PDB ID 9B8S (2)) was used as the initial model. The structures of POLE1-NTD and human PCNA were manually docked into the EM maps using UCSF ChimeraX (11). The models were manually adjusted and rebuilt in Coot(8, 12) to fit the EM maps. The original maps were sharpened by EMReady(13) to facilitate modeling of the T/P in the mismatch editing state. The P/T DNA was modified from our published structure in the pol state (PDB ID 9B8T(2)) to fit the T/P EM density in the three proofreading states. Only bases near the mismatched primer 3′-end were rebuilt. Unresolved DNA regions were omitted in all the states. The manually built initial models were subjected to several iterations of real-space refinement in PHENIX (14) and further manual adjustment in Coot (12). All final atomic models were validated using MolProbity (15). The 3D reconstruction and model refinement statistics are provided in **Supplementary Table 1**. Structural figures were prepared in the UCSF ChimeraX(11).

### Data Availability

Two cryo-EM 3D maps (3.88 and 3.81 Å) of the ternary human Polε-core– PCNA–T49/P35 (pre-existing mismatch) in the blocked conformations, along with their corresponding atomic models, have been deposited in the Electron Microscopy Data Bank (https://www.ebi.ac.uk/pdbe/emdb/) and the Protein Data Bank (https://www.rcsb.org) under the following accession codes: EMD-49302 and 9NE9 (3.88 Å, The blocked conformation I) and EMD-49303 and 9NEA (3.81 Å, The blocked conformation II). The three 3D EM maps of human Polε-core–PCNA–T47/P29 (with in situ generated mismatch) in three proofreading states (3.60 Å, 3.53 Å, 3.11Å) have been deposited in the EMDB and PDB with accession codes: EMD-49301 and 9NE8 (3.60 Å, Mismatch-locking state), EMD-49300 and 9NE7 (3.53 Å, Pol-backtracking state), and EMD-49299 and 9NE6 (3.11 Å, Mismatch-editing state).

**SI Appendix, Video S1. A detailed depiction of the DNA proofreading process by the Polε– PCNA holoenzyme.** Morphing of Polε–PCNA proofreading intermediate structures to show the DNA translocation steps and the coordinated conformational changes of Polε.

## REFERENCES

1. A. Kornberg, DNA replication. Trends Biochem. Sci. 9, 122–124 (1984).

2. L. D. Langston, M. O’Donnell, DNA replication: keep moving and don’t mind the gap. Mol. Cell 23, 155–160 (2006).

3. P. M. Burgers, T. A. Kunkel, Eukaryotic DNA replication fork. Annu. Rev. Biochem. 86, 417–438 (2017).

4. A. Johnson, M. O’Donnell, Cellular DNA replicases: components and dynamics at the replication fork. Annu. Rev. Biochem. 74, 283–315 (2005).

5. Z. F. Pursell, I. Isoz, E.-B. Lundström, E. Johansson, T. A. Kunkel, Yeast DNA polymerase ε participates in leading-strand DNA replication. Science 317, 127–130 (2007).

6. E. Johansson, N. Dixon, Replicative DNA polymerases. Cold Spring Harb. perspect. biol. 5, a012799 (2013).

7. T. A. Kunkel, P. M. Burgers, Arranging eukaryotic nuclear DNA polymerases for replication: Specific interactions with accessory proteins arrange Pols α, δ, and ɛ in the replisome for leading-strand and lagging-strand DNA replication. Bioessays 39, 1700070 (2017).

8. A. Bernad, L. Blanco, J. Lázaro, G. Martin, M. Salas, A conserved 3′→ 5′ exonuclease active site in prokaryotic and eukaryotic DNA polymerases. Cell 59, 219–228 (1989).

9. T. A. Steitz, DNA polymerases: structural diversity and common mechanisms. J. Biol. Chem. 274, 17395–17398 (1999).

10. A. Bębenek, I. Ziuzia-Graczyk, Fidelity of DNA replication—a matter of proofreading. Curr. Genet. 64, 985–996 (2018).

11. O. Chilkova, B.-H. Jonsson, E. Johansson, The quaternary structure of DNA polymerase ε from Saccharomyces cerevisiae. J. Biol. Chem. 278, 14082–14086 (2003).

12. Z. F. Pursell, T. A. Kunkel, DNA polymerase ε: a polymerase of unusual size (and complexity). Proc. Natl. Acad. Sci. U.S.A. 82, 101–145 (2008).

13. T. H. Tahirov, K. S. Makarova, I. B. Rogozin, Y. I. Pavlov, E. V. Koonin, Evolution of DNA polymerases: an inactivated polymerase-exonuclease module in Pol ε and a chimeric origin of eukaryotic polymerases from two classes of archaeal ancestors. Bio. Direct 4, 1–11 (2009).

14. D. Kazlauskas, M. Krupovic, J. Guglielmini, P. Forterre, Č. Venclovas, Diversity and evolution of B-family DNA polymerases. Nucleic acids res. 48, 10142–10156 (2020).

15. Z. Yuan, R. Georgescu, G. D. Schauer, M. E. O’Donnell, H. Li, Structure of the polymerase ε holoenzyme and atomic model of the leading strand replisome. Nat. Commun. 11, 3156 (2020).

16. J. C. Zhou et al., CMG–Pol epsilon dynamics suggests a mechanism for the establishment of leading-strand synthesis in the eukaryotic replisome. Proc. Natl. Acad. Sci. U.S.A. 114, 4141–4146 (2017).

17. I. Isoz, U. Persson, K. Volkov, E. Johansson, The C-terminus of Dpb2 is required for interaction with Pol2 and for cell viability. Nucleic acids res. 40, 11545–11553 (2012).

18. L. D. Langston et al., CMG helicase and DNA polymerase ε form a functional 15-subunit holoenzyme for eukaryotic leading-strand DNA replication. Proc. Natl. Acad. Sci. U.S.A. 111, 15390–15395 (2014).

19. P. Goswami et al., Structure of DNA-CMG-Pol epsilon elucidates the roles of the non-catalytic polymerase modules in the eukaryotic replisome. Nat. Commun. 9, 5061 (2018).

20. Z. Xu et al., Synergism between CMG helicase and leading strand DNA polymerase at replication fork. Nat. Commun. 14, 5849 (2023).

21. M. L. Jones, Y. Baris, M. R. Taylor, J. T. Yeeles, Structure of a human replisome shows the organisation and interactions of a DNA replication machine. EMBO J. 40, e108819 (2021).

22. F. J. Asturias et al., Structure of Saccharomyces cerevisiae DNA polymerase epsilon by cryo–electron microscopy. Nat. Struct. Mol. Biol. 13, 35–43 (2006).

23. M. Hogg et al., Structural basis for processive DNA synthesis by yeast DNA polymerase ɛ. Nat. Struct. Mol. Biol. 21, 49–55 (2014).

24. Q. He, F. Wang, N. Y. Yao, M. E. O’Donnell, H. Li, Structures of the human leading strand Polε–PCNA holoenzyme. Nat. Commun. 15, 7847 (2024).

25. J. J. Roske, J. T. P. Yeeles, Structural basis for processive daughter-strand synthesis and proofreading by the human leading-strand DNA polymerase Pol ε. Nat. Struct. Mol. Biol. 31, 1921–1931 (2024).

26. N. Singh, E. Johansson, Clamping Pol ε to the leading strand. Nat. Struct. Mol. Biol. 31, 1–2 (2024).

27. O. Chilkova et al., The eukaryotic leading and lagging strand DNA polymerases are loaded onto primer-ends via separate mechanisms but have comparable processivity in the presence of PCNA. Nucleic acids res. 35, 6588–6597 (2007).

28. J. T. Yeeles, A. Janska, A. Early, J. F. Diffley, How the eukaryotic replisome achieves rapid and efficient DNA replication. Mol. Cell 65, 105–116 (2017).

29. E. M. Boehm, M. S. Gildenberg, M. T. Washington, The many roles of PCNA in eukaryotic DNA replication. The Enzymes 39, 231–254 (2016).

30. T. Mondol, J. L. Stodola, R. Galletto, P. M. Burgers, PCNA accelerates the nucleotide incorporation rate by DNA polymerase δ. Nucleic acids res. 47, 1977–1986 (2019).

31. F. Zheng, R. Georgescu, N. Y. Yao, H. Li, M. E. O’Donnell, Cryo-EM structures reveal that RFC recognizes both the 3′-and 5′-DNA ends to load PCNA onto gaps for DNA repair. eLife 11, e77469 (2022).

32. X. Liu, C. Gaubitz, J. Pajak, B. A. Kelch, A second DNA binding site on RFC facilitates clamp loading at gapped or nicked DNA. Elife 11, e77483 (2022).

33. M. Schrecker et al., Multistep loading of a DNA sliding clamp onto DNA by replication factor C. eLife 11, e78253 (2022).

34. Z. Yuan et al., Mechanism of PCNA loading by Ctf18-RFC for leading-strand DNA synthesis. Science 385, eadk5901 (2024).

35. R. Lamichhane, S. Y. Berezhna, E. Van, D. P. Millar, Dynamics of site switching in DNA polymerase. Biophys. J. 104, 368a (2013).

36. L. Xu, M. T. Halma, G. J. Wuite, Mapping fast DNA polymerase exchange during replication. Nat. Commun. 15, 5328 (2024).

37. L. S. Beese, V. Derbyshire, T. A. Steitz, Structure of DNA polymerase I Klenow fragment bound to duplex DNA. Science 260, 352–355 (1993).

38. T. Dodd et al., Polymerization and editing modes of a high-fidelity DNA polymerase are linked by a well-defined path. Nat. Commun. 11, 5379 (2020).

39. M. Wang et al., Insights into base selectivity from the 1.8 Å resolution structure of an RB69 DNA polymerase ternary complex. Biochemistry 50, 581–590 (2011).

40. M. Hogg, S. S. Wallace, S. Doublié, Crystallographic snapshots of a replicative DNA polymerase encountering an abasic site. EMBO J. 23, 1483–1493 (2004).

41. Y. Shamoo, T. A. Steitz, Building a replisome from interacting pieces: sliding clamp complexed to a peptide from DNA polymerase and a polymerase editing complex. Cell 99, 155–166 (1999).

42. J. Gouge, C. Ralec, G. Henneke, M. Delarue, Molecular recognition of canonical and deaminated bases by P. abyssi family B DNA polymerase. J. Mol. Biol. 423, 315–336 (2012).

43. J. Park, G. K. Herrmann, P. G. Mitchell, M. B. Sherman, Y. W. Yin, Polγ coordinates DNA synthesis and proofreading to ensure mitochondrial genome integrity. Nat. Struct. Mol. Biol. 30, 812–823 (2023).

44. G. Buchel et al., Structural basis for DNA proofreading. Nat. Commun. 14, 8501 (2023).

45. S. T. Lovett, The DNA exonucleases of Escherichia coli. EcoSal Plus 4, 10.1128/ecosalplus. 1124.1124. 1127 (2011).

46. J. Wang, P. Yu, T. Lin, W. Konigsberg, T. Steitz, Crystal structures of an NH2-terminal fragment of T4 DNA polymerase and its complexes with single-stranded DNA and with divalent metal ions. Biochemistry 35, 8110–8119 (1996).

47. R. Fernandez-Leiro et al., Self-correcting mismatches during high-fidelity DNA replication. Nat. Struct. Mol. Biol. 24, 140–143 (2017).

48. L. Betancurt-Anzola et al., Molecular basis for proofreading by the unique exonuclease domain of Family-D DNA polymerases. Nat. Commun. 14, 8306 (2023).

49. C. R. Bulock, X. Xing, P. V. Shcherbakova, Mismatch repair and DNA polymerase δ proofreading prevent catastrophic accumulation of leading strand errors in cells expressing a cancer-associated DNA polymerase ɛ variant. Nucleic acids res. 48, 9124–9134 (2020).

50. E. Shinbrot et al., Exonuclease mutations in DNA polymerase epsilon reveal replication strand specific mutation patterns and human origins of replication. Genome Res. 24, 1740–1750 (2014).

51. R. A. Ganai, G. O. Bylund, E. Johansson, Switching between polymerase and exonuclease sites in DNA polymerase ε. Nucleic acids res. 43, 932–942 (2015).

52. S. J. Johnson, L. S. Beese, Structures of mismatch replication errors observed in a DNA polymerase. Cell 116, 803–816 (2004).

53. S. Xia, W. H. Konigsberg, Mispairs with Watson-Crick base-pair geometry observed in ternary complexes of an RB69 DNA polymerase variant. Protein Sci. 23, 508–513 (2014).

54. S. Xia, J. Wang, W. H. Konigsberg, DNA mismatch synthesis complexes provide insights into base selectivity of a B family DNA polymerase. J. Am. Chem. Soc. 135, 193–202 (2013).

55. M. Hogg, P. Aller, W. Konigsberg, S. S. Wallace, S. Doublié, Structural and biochemical investigation of the role in proofreading of a β hairpin loop found in the exonuclease domain of a replicative DNA polymerase of the B family. J. Biol. Chem. 282, 1432–1444 (2007).

56. T. L. Dangerfield, S. Kirmizialtin, K. A. Johnson, Substrate specificity and proposed structure of the proofreading complex of T7 DNA polymerase. J. Biol. Chem. 298 (2022).

57. A. Del Prado et al., Noncatalytic aspartate at the exonuclease domain of proofreading DNA polymerases regulates both degradative and synthetic activities. Proc. Natl. Acad. Sci. U.S.A. 115, E2921–E2929 (2018).

58. A. Trzemecka, D. Płochocka, A. Bebenek, Different behaviors in vivo of mutations in the β hairpin loop of the DNA polymerases of the closely related phages T4 and RB69. J. Mol. Biol. 389, 797–807 (2009).

59. M. K. Swan, R. E. Johnson, L. Prakash, S. Prakash, A. K. Aggarwal, Structural basis of high-fidelity DNA synthesis by yeast DNA polymerase δ. Nat. Struct. Mol. Biol. 16, 979–986 (2009).

60. V. Parkash et al., Structural consequence of the most frequently recurring cancer-associated substitution in DNA polymerase ε. Nat. Commun. 10, 373 (2019).

61. D. P. Kane, P. V. Shcherbakova, A common cancer-associated DNA polymerase ɛ mutation causes an exceptionally strong mutator phenotype, indicating fidelity defects distinct from loss of proofreading. Cancer Res. 74, 1895–1901 (2014).

62. R. Fujisawa, E. Ohashi, K. Hirota, T. Tsurimoto, Human CTF18-RFC clamp-loader complexed with non-synthesising DNA polymerase ε efficiently loads the PCNA sliding clamp. Nucleic Acids Res. 45, 4550–4563 (2017).

63. K. Stokes, A. Winczura, B. Song, G. De Piccoli, D. B. Grabarczyk, Ctf18-RFC and DNA Pol ɛ form a stable leading strand polymerase/clamp loader complex required for normal and perturbed DNA replication. Nucleic acids res. 48, 8128–8145 (2020).

64. Q. He, F. Wang, M. E. O’Donnell, H. Li, Cryo-EM reveals a nearly complete PCNA loading process and unique features of the human alternative clamp loader CTF18-RFC. Proc. Natl. Acad. Sci. U.S.A. 121, e2319727121 (2024).

65. K. Bebenek, J. Abbotts, S. Wilson, T. Kunkel, Error-prone polymerization by HIV-1 reverse transcriptase. Contribution of template-primer misalignment, miscoding, and termination probability to mutational hot spots. J. Biol. Chem. 268, 10324–10334 (1993).

66. S. Fujii et al., DNA replication errors produced by the replicative apparatus of Escherichia coli. J. Mol. Biol. 289, 835–850 (1999).

67. D. Canceill, E. Viguera, S. D. Ehrlich, Replication slippage of different DNA polymerases is inversely related to their strand displacement efficiency. J. Biol. Chem. 274, 27481–27490 (1999).

68. E. Heitzer, I. Tomlinson, Replicative DNA polymerase mutations in cancer. Curr. Opin. Genet. Dev. 24, 107–113 (2014).

69. S. R. Barbari, D. P. Kane, E. A. Moore, P. V. Shcherbakova, Functional analysis of cancer-associated DNA polymerase ε variants in Saccharomyces cerevisiae. G3: Genes Genomes Genet. 8, 1019-1029 (2018).

70. F. Wang, Q. He, H. Li, Human polymerase epsilon bound to PCNA and DNA with a pre-existing mismatch in the blocked conformation I. Electron Microscopy Data Bank. https://www.ebi.ac.uk/emdb/EMD-49302. Deposited 19 February 2025.

71. F. Wang, Q. He, H. Li, Human polymerase epsilon bound to PCNA and DNA with a pre-existing mismatch in the blocked conformation I. Protein Data Bank. https://www.rcsb.org/structure/9NE9. Deposited 19 February 2025.

72. F. Wang, Q. He, H. Li, Human polymerase epsilon bound to PCNA and DNA with a pre-existing mismatch in the blocked conformation II. Electron Microscopy Data Bank. https://www.ebi.ac.uk/emdb/EMD-49303. Deposited 19 February 2025.

73. F. Wang, Q. He, H. Li, Human polymerase epsilon bound to PCNA and DNA with a pre-existing mismatch in the blocked conformation II. Protein Data Bank. https://www.rcsb.org/structure/9NEA. Deposited 19 February 2025.

74. F. Wang, Q. He, H. Li, Human polymerase epsilon bound to PCNA and DNA with an in-situ-generated mismatch in the mismatch-locking state. Electron Microscopy Data Bank. https://www.ebi.ac.uk/emdb/EMD-49301. Deposited 19 February 2025.

75. F. Wang, Q. He, H. Li, Human polymerase epsilon bound to PCNA and DNA with an in-situ-generated mismatch in the mismatch-locking state. Protein Data Bank. https://www.rcsb.org/structure/9NE8. Deposited 19 February 2025.

76. F. Wang, Q. He, H. Li, Human polymerase epsilon bound to PCNA and DNA with an in-situ-generated mismatch in the Pol-backtracking state state. Electron Microscopy Data Bank. https://www.ebi.ac.uk/emdb/EMD-49300. Deposited 19 February 2025.

77. F. Wang, Q. He, H. Li, Human polymerase epsilon bound to PCNA and DNA with an in-situ-generated mismatch in the Pol-backtracking state state. Protein Data Bank. https://www.rcsb.org/structure/9NE7. Deposited 19 February 2025.

78. F. Wang, Q. He, H. Li, Human polymerase epsilon bound to PCNA and DNA with an in-situ-generated mismatch in the mismatch-editing state. Electron Microscopy Data Bank. https://www.ebi.ac.uk/emdb/EMD-49299. Deposited 19 February 2025.

79. F. Wang, Q. He, H. Li, Human polymerase epsilon bound to PCNA and DNA with an in-situ-generated mismatch in the mismatch-editing state. Protein Data Bank. https://www.rcsb.org/structure/9NE6. Deposited 19 February 2025.

## References for SI Appendix

1. F. Weissmann et al., biGBac enables rapid gene assembly for the expression of large multisubunit protein complexes. Proc. Natl. Acad. Sci. U.S.A. 113, E2564–E2569 (2016).

2. Q. He, F. Wang, N. Y. Yao, M. E. O’Donnell, H. Li, Structures of the human leading strand Polε–PCNA holoenzyme. Nat. Commun. 15, 7847 (2024).

3. F. Wang, Q. He, N. Y. Yao, M. E. O’Donnell, H. Li, The human ATAD5 has evolved unique structural elements to function exclusively as a PCNA unloader. Nat. Struct. Mol. Biol. 31, 1680–1691 (2024).

4. J. J. Roske, J. T. P. Yeeles, Structural basis for processive daughter-strand synthesis and proofreading by the human leading-strand DNA polymerase Pol ε. Nat. Struct. Mol. Biol. 31, 1921–1931 (2024)

5. D. N. Mastronarde, Automated electron microscope tomography using robust prediction of specimen movements. J. Struct. Biol. 152, 36–51 (2005).

6. S. Q. Zheng et al., MotionCor2: anisotropic correction of beam-induced motion for improved cryo-electron microscopy. Nat. methods 14, 331–332 (2017).

7. A. Punjani, J. L. Rubinstein, D. J. Fleet, M. A. Brubaker, cryoSPARC: algorithms for rapid unsupervised cryo-EM structure determination. Nat. methods 14, 290–296 (2017).

8. A. Punjani, D. J. Fleet, 3D variability analysis: Resolving continuous flexibility and discrete heterogeneity from single particle cryo-EM. J. Struct. Biol. 213, 107702 (2021).

9. D. Kimanius et al., Data-driven regularization lowers the size barrier of cryo-EM structure determination. Nat. Methods 21, 1216–1221 (2024).

10. A. Burt et al., An image processing pipeline for electron cryo-tomography in RELION-5. FEBS Open Bio (2024).

11. E. F. Pettersen et al., UCSF ChimeraX: Structure visualization for researchers, educators, and developers. Protein Sci. 30, 70–82 (2021).

12. P. Emsley, B. Lohkamp, W. G. Scott, K. Cowtan, Features and development of Coot. Acta Crystallogr. D 66, 486–501 (2010).

13. J. He, T. Li, S.-Y. Huang, Improvement of cryo-EM maps by simultaneous local and non-local deep learning. Nat. Commun. 14, 3217 (2023).

14. P. V. Afonine et al., Real-space refinement in PHENIX for cryo-EM and crystallography. Acta Crystallogr. D 74, 531–544 (2018).

15. C. J. Williams et al., MolProbity: More and better reference data for improved all-atom structure validation. Protein Sci. 27, 293–315 (2018).

